# Breeding cassava for intercropping with cowpea: monoculture selection captures most intercrop selection gain, but targeted testing remains necessary

**DOI:** 10.64898/2026.08.31.748306

**Authors:** Oluwaseye Gideon Oyebode, Ayankami Toye, Iluebbey Peter, Ukoabasi Okon Ekanem, Ogwuche O. ThankGod, Ismail Yusuf Rabbi, Marnin Wolfe

**Affiliations:** Cassava breeding unit, International Institute of Tropical Agriculture (IITA), Ibadan, Nigeria; Wolfe Lab, Department of Crop, Soils, and Environmental Sciences, Auburn University, AL, USA

**Keywords:** Cassava, Cowpea, Intercrop, Testers, General and specific mixing ability, Producer and associate effects, Selection efficiency, Genetic correlation, Genotype × management interaction

## Abstract

Intercropping dominates smallholder cassava production in sub-Saharan Africa, yet cassava breeding programs evaluate genotypes exclusively under monoculture. Despite consistently reported system-level yield advantages, cassava yield is reduced by 17–51% under intercropping, indicating a need to reduce this competitive disadvantage through breeding. However, the quantitative-genetic foundations of intercrop breeding remain uncharacterized for tropical root-crop systems. We hypothesized that monoculture selection would capture most, but not all, genetic merit for intercropping and that a limited tester set would be sufficient if general mixing ability predominated.

We evaluated 120 cassava clones previously selected under monoculture in IITA advanced yield trials, testing them under monoculture and in intercrop with two contrasting cowpea varieties across two years at Ibadan, Nigeria. Spatial mixed models quantified genetic variation, genotype-by-cropping-system interaction, cross-system genetic relationships, realized selection gain, mixing ability, tester effects, and land equivalent ratio.

Intercropping reduced cassava fresh root yield by 19%, but total land equivalent ratios exceeded 1.0 for all clones, confirming a system-level land-use advantage. Cassava performance under intercropping was heritable, with estimates of 0.50–0.75, and genotype-by-cropping-system interaction was not significant. Genetic correlations between monoculture and intercrop performance were high and approached unity (rg = 0.92–0.99), and selection efficiency was 26–44% at 10% intensity, confirming that high genetic correlation does not guarantee effective indirect selection. Monoculture selection captured approximately two-thirds of direct intercrop gain. General mixing ability dominated, specific mixing ability was negligible, and producer effects explained 20–47% of intercrop variance. The two architecturally and phenologically contrasting cowpea varieties had limited influence on cassava rankings. Here, we show for the first time that cassava breeding for intercropping can retain monoculture selection during early stages while adding two representative cowpea testers at the advanced-trial stage. This staged strategy aligns cassava breeding with diversified smallholder systems without creating a separate pipeline.

## 1 Introduction

Intercropping is widespread in cassava production, with an estimated 30–50% of cassava produced globally grown in intercrop systems (Parmar et al. 2017; Weerarathne et al. 2017), and it has re- emerged as an important strategy for sustainable intensification (Adam et al., 2025; Sabatier et al. 2026). Compared with monoculture, intercropping can substantially increase overall productivity and land-use efficiency, with approximately 20–30% productivity advantage over monoculture, together with greater yield stability and more efficient use of soil moisture and nutrients (Raseduzzaman and Jensen 2017; Li et al. 2023; Jensen et al. 2020). In sub-Saharan Africa, where production is predominantly rainfed and 70– 83% of smallholder farmers practice intercropping, it contributes to both crop diversification and climate risk management (Himmelstein et al. 2017; Martin-Guay et al. 2018; Gmakouba et al. 2024). Meta-analyses report an average land equivalent ratio (LER) of approximately 1.45 in the region, with root– legume systems performing particularly well under variable moisture conditions (Himmelstein et al. 2017; Adam et al. 2025). Cassava (*Manihot esculenta* Crantz), the most important tropical root crop (Parmar et al. 2017), is commonly intercropped with legumes in sub-Saharan Africa, where cassava– legume systems represent the most promising climate-resilient diversification strategy (Dettweiler et al. 2023; Adam et al. 2025).

Despite the favorable land-use efficiency of cassava–legume intercropping, cassava root yield is frequently reduced by 17–51% relative to monoculture (Dapaah et al. 2003; Hidoto and Loha 2013; Nyi et al. 2014; Mwebaze et al. 2024). The contrast between increased total-system productivity and reduced cassava yield suggests that further improvement of these systems may require breeding to reduce cassava’s competitive disadvantage, alongside agronomic management (Cenpukdee and Fukai 1992c; Brooker et al. 2015; Annicchiarico et al. 2021; Haug and Bourke 2025). Cassava breeding programs, however, have historically evaluated and selected genotypes almost exclusively under monoculture (Dettweiler et al. 2023). This creates a mismatch between the breeding environment and the diversified production systems in which many cassava varieties are grown (Rubiales et al. 2023; Hohmann et al. 2026). Traits that improve performance in intercrop may not be functional under monoculture evaluation, whereas highly competitive varieties selected in monoculture may suppress the companion crop and reduce total-system performance (Annicchiarico et al. 2019; Abou-Khater et al. 2024; Hohmann et al. 2026). This raises a fundamental question: does monoculture selection adequately capture genetic potential for intercrop performance, or does cassava–cowpea intercropping require direct evaluation under intercropping? Early cassava studies anticipated this question: Cenpukdee and Fukai (1992a, b) showed that the suitability of monoculture selection for intercrop breeding depended on the competitive ability of the companion legume, with sole-crop selection more applicable when cassava was paired with a less competitive associate.

The quantitative-genetic foundations of breeding for intercropping systems were formalized by Wright (1985), who partitioned intercrop performance into producer and associate effects. Producer effects describe a genotype’s capacity to maintain its own performance in intercrop, whereas associate effects describe its influence on the performance of the companion crop (Haug et al. 2021, 2023). Their combination sums to general mixing ability (GMA), which represents performance across partners, while specific mixing ability (SMA) captures effects unique to particular genotype combinations (Wright 1985; Sampoux et al. 2020; Haug et al. 2023). This distinction has direct implications for breeding strategy.

When GMA predominates and genotype rankings remain reasonably stable across partners, with a non-significant genotype by management interaction, broadly compatible genotypes may be identified using a limited set of testers (Moore et al. 2022; Haug et al. 2023). Conversely, substantial SMA would require partner-specific selection and a larger, more costly testing system (Haug et al. 2021, 2023; Sampoux et al. 2020).

Whether direct intercrop evaluation is required also depends on the genetic relationship between monoculture and intercrop performance. Selection in monoculture can generate a correlated response under intercropping when monoculture performance is positively associated with the genetic effects governing intercrop performance (Sampoux et al. 2020). Simulation evidence further indicates that the optimal stage for introducing direct intercrop testing depends jointly on trait heritability and the genetic correlation between monoculture and intercrop performance, with higher correlations generally allowing intercrop evaluation to be delayed to later breeding stages (Dubey et al. 2024). In temperate intercrops, indirect selection based on monoculture performance has achieved approximately 52–64% of the efficiency of direct intercrop selection, depending partly on the genetic correlation between systems (Annicchiarico et al. 2021). However, a high genetic correlation does not necessarily ensure agreement among selected genotypes from monoculture and intercrop performance. In common bean–maize intercrops, Zimmermann (1996) reported moderate-to-high correlations between monoculture and intercrop yield, while selection coincidence frequently remained below 50% because of genotype re-ranking. Therefore, the usefulness of monoculture selection depends not only on genetic correlation but also on whether it identifies the same superior genotypes and delivers comparable selection gains under intercropping (Annicchiarico et al. 2019).

A major operational constraint in intercrop breeding is the intractable number of possible cassava × cowpea combinations that could be evaluated (Haug et al. 2021, 2023). A limited set of companion genotypes, termed testers by analogy with hybrid breeding, can instead be used to reveal the mixing ability of the species in the intercrop (Moutier et al. 2022). Evidence from forage and cereal–legume mixtures indicate that GMA often exceeds SMA and that carefully chosen testers or incomplete-factorial designs can substantially reduce the testing burden (Holland and Brummer 1999; Annicchiarico et al. 2019; Haug et al. 2023). This operational simplification is particularly relevant for resource-constrained breeding programs. Thus, targeted intercrop evaluation could potentially be embedded within an existing breeding pipeline rather than requiring a parallel testing program (Dubey et al. 2024; Hohmann et al. 2026).

Most quantitative-genetic studies of intercrop breeding have focused on temperate cereal–legume systems, whereas tropical cassava-based systems remain poorly characterized. This gap is critical because cassava differs fundamentally from annual cereals in growth duration, architecture, phenology, and biomass allocation, characteristics that may alter genetic relationships between monoculture and intercrop performance. Cassava fresh-root yield and cowpea grain yield also differ strongly in scale, creating a risk that naïve aggregation will allow one crop to dominate a combined response and distort estimates of intercrop value and mixing ability (Wright 1985).

Collectively, these gaps leave several key questions unanswered: the extent to which monoculture performance predicts intercrop performance, how genetic effects are partitioned between cassava and cowpea partners, if cowpea tester identity alters cassava selection decisions, and whether intercropping environments retain sufficient exploitable genetic variation for effective selection (Bourke et al. 2021; Moore et al. 2022). To address these gaps, the objectives of this study were to: (1) quantify exploitable genetic variation under cassava–cowpea intercropping, evaluate genotype-by-cropping-system interaction, and characterize system productivity; (2) determine the extent to which monoculture selection predicts intercrop performance and realized selection gain; and (3) partition intercrop performance into producer, associate, general mixing, and specific mixing effects and determine whether cowpea tester choice influences cassava selection.

## 2 Materials and methods

### 2.1 Field trial, planting arrangement and experimental design

Cassava–cowpea intercropping trials were conducted over two seasons (August 2023 to August 2024, and September 2024 to October 2025) at the International Institute of Tropical Agriculture (IITA), Ibadan, Nigeria (7°24′N, 3°54′E; ∼230 m a.s.l.), on rainfed Ferric Luvisols of sandy-loam surface texture (Fig. 1). The site has a sub-humid tropical climate; the two seasons had comparable thermal time and total rainfall (∼1,750 mm) but differed in distribution, with a pronounced December–January dry spell in year 1 and a shorter low-rainfall period in year 2 (Fig. S1).

**Fig. 1.**
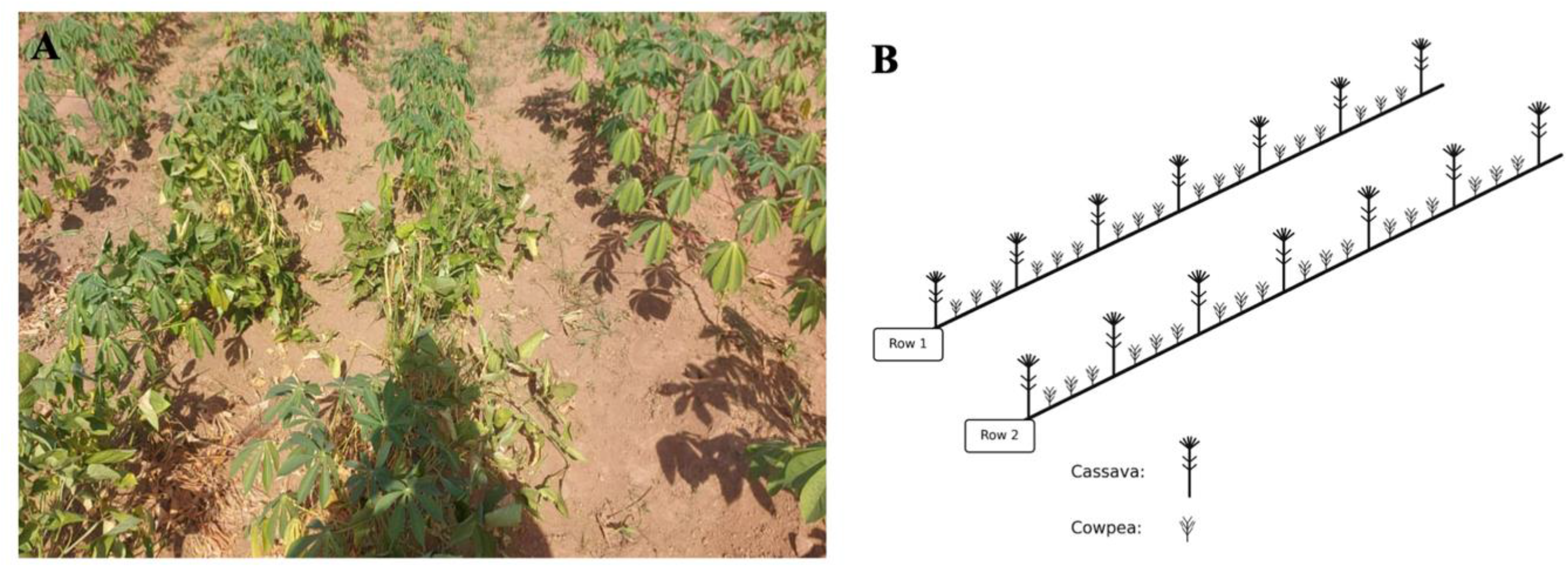
The cassava–cowpea intercropping system used in this study. (a) Plot view of the intercrop at IITA, Ibadan, Nigeria, showing cassava plants with cowpea interplanted between them within the rows. (b) Schematic of the planting arrangement: each plot comprised two cassava rows (5 × 3 m; 15 m²), with cassava spaced at 1.0 m between rows and 0.8 m within rows, and cowpea interplanted within the cassava rows at 0.25 m in an additive design (three cowpea plants between consecutive cassava stands). Photo: Oluwaseye G. Oyebode.

The experiment used a row–column design with two replicates per year, monoculture and intercrop treatments randomized within replicates. Plots were 5 × 3 m (15 m²), each with two cassava rows at 1.0 × 0.8 m (∼12,500 plants ha⁻¹). Cowpea was interplanted within cassava rows at 0.25 m in this additive design, and grown as monoculture at 1.0 × 0.25 m. Each year comprised 362 treatments (120 cassava monocultures, 240 cassava × two-tester combinations, and 2 cowpea monocultures) in two replicates, giving 1,448 plots across the two seasons. Agronomic management and practices for cassava and cowpea followed standard IITA practices (Rabbi et al. 2022; Popoola et al. 2024).

### 2.2 Cassava genotypes and cowpea testers

120 cassava clones (also referred to as genotypes in modeling equations) were selected from the IITA breeding population archived in Cassavabase (www.cassavabase.org), originating from Advanced Yield Trials (Rabbi et al. 2022). Multi-environment mixed-model analyses followed by k-means clustering were used to select clones representative of the breeding population’s genetic diversity. Two farmer-preferred cowpea varieties with contrasting maturity and architecture traits linked to legume competitive ability in intercrops (Annicchiarico and Piano 1994; Wang et al. 2004, 2006) were selected as testers: Oloyin, a late-maturing variety with a spreading growth habit, and IT08K-150-12 (SAMPEA 19), an elite IITA line with early maturity and an erect growth habit (Omoigui et al. 2018).

### 2.3 Trait measurement

Cassava plots were harvested 12 months after planting and cowpea plots at maturity. For cassava, fresh root yield (kg plot⁻¹), root dry matter content (%), harvest index, and the number of harvested plants were recorded; for cowpea, grain yield (g plot⁻¹) and the number of harvested plants were recorded. The numbers of harvested cassava and cowpea plants were used as covariates (Section 2.6), to adjust genetic estimates for plot-level establishment differences.

### 2.4 Intercrop productivity metrics and the yield-scale asymmetry

Cassava fresh root yield and cowpea grain yield differed markedly in numerical scale. To assess whether this asymmetry affected the determination of intercrop merit, six metrics were compared: the sum of yields in their native units, the sum after conversion to a common mass unit (kg), an economic-value index based on crop-specific market prices, the first principal component (PC1) of standardized component yields, a partial-land-equivalent-ratio-based index, and a z-score index. For the economic metric, cassava fresh root weight was valued at ₦80 kg⁻¹, Oloyin cowpea at ₦1500 kg⁻¹, and IT08K-150- 12 at ₦1300 kg⁻¹, reflecting current market prices in the trial area at the time of analysis.

For the z-score metric, cassava fresh root yield and cowpea grain yield were standardized to zero mean and unit variance within year and summed so that the two component crops contributed on comparable scales. This formulation corresponds to Wright’s (1985) selection-index approach with variance- equalizing weights inversely proportional to the standard deviation of each component. PC1 was derived from the two standardized component yields, whereas the pLER-based metric expressed cassava yield relative to the corresponding clone monoculture and cowpea grain yield relative to the monoculture yield of the same tester.

Scale dependence among metrics was evaluated using the cassava-cowpea contribution ratio, decomposition of composite variance into cassava, cowpea, and covariance components, and Pearson correlations between each composite metric and the two component yields. Based on these comparisons, the z-score composite was retained as the intercrop-merit phenotype for all subsequent mixing-ability analyses, including estimation of GMA, SMA, and producer and associate effects (Section 2.11).

### 2.5 Statistical analysis

Analyses were conducted in R with ASReml-R v4.2 (Butler et al. 2018), fitting linear mixed models with variance components estimated by residual maximum likelihood (REML), both across and within years. Field heterogeneity was modelled throughout with a separable first-order autoregressive (AR1 × AR1) residual structure over row and column positions. Monoculture and intercrop datasets were first analyzed separately to estimate system-specific genetic parameters.

### 2.6 Mixed models for monoculture and intercrop

Across years, the following model was fitted, with the cowpea-density covariate (*β*_2_*P_ijk_*) included for intercrop only:

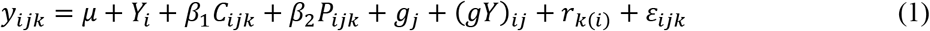

where *y_ijk_* is the plot observation, *μ* the overall mean, *Y_i_* the fixed year effect, *C_ijk_* and *P_ijk_* the numbers of harvested cassava and cowpea plants (centred within year), 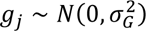 the random clone effect, 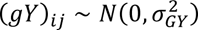 the clone × year interaction, *r_k_*_(_*_i_*_)_ the replicate within year, and 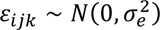 the spatially correlated residual. Within-year analyses excluded the year and clone × year terms. Entry-mean broad-sense heritability across years was estimated as:

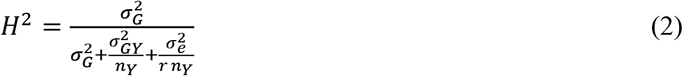

with *n_Y_* = 2 years and *r* = 2 replicates per year, estimated separately for each system. For within-year analyses this reduced to 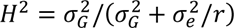

### 2.7 Cassava genotype × management interaction

A combined model across years tested whether clone performance differed between systems:

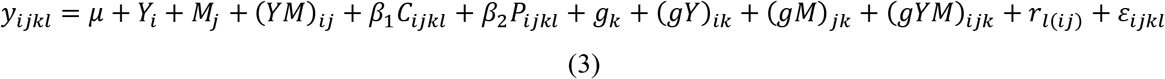

where *M_j_* is the fixed management effect (monoculture or intercrop) and (*YM*)*_ij_* the year × management interaction; 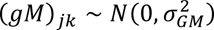 is the clone × management interaction and 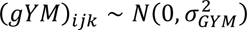 the three-way interaction. Other terms are as in Eq. 1. Within-year analyses omitted year-related terms. The variance components 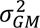, 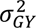 and 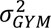 were tested by likelihood-ratio tests against reduced models.

### 2.8 Genetic correlation between monoculture and intercrop

Genetic correlation was estimated with a bivariate model stacking both systems into one response, with system (monoculture, intercrop) as a two-level trait factor. Fixed effects were system, system × year, system × tester, and a system × stand-count covariate centered within system and year. The between-system genetic effect was unstructured, estimating the two genetic variances and their covariance, and the genetic correlation was:

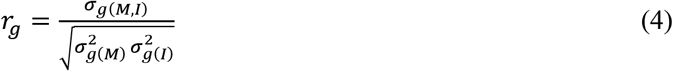

Two model features were selected empirically rather than assumed. Because monoculture and intercrop plots occupied unique positions on one physical field, a common within-year AR1 × AR1 grid on the original coordinates was compared with separate per-system grids; and a diagonal genotype-by-year structure (system-specific year variances, covariance fixed at zero) was compared with an unstructured form estimating the across-system year covariance. The four models were compared by AIC, BIC, and likelihood-ratio tests where nested (Table S1). A common grid with a diagonal genotype-by-year structure was retained as the primary model for all three traits, on grounds of convergence reliability and consistency across traits; the unstructured form is reported as a sensitivity analysis in the supplementary material (Table S1).

The hypothesis of no genetic differentiation between systems (*H*_0_: *r_g_* = 1) was tested by comparing the unstructured model with a constrained rank-one, fixed-unit-correlation model that removes exactly one free parameter. Because *r_g_* = 1 lies on the parameter-space boundary, the statistic was referred to a 0.5:0.5 mixture of *χ*^2^(0) and *χ*^2^(1) (Self and Liang 1987). Pearson and Spearman correlations between clone BLUPs from the system-specific fits of Eq. 1 (monoculture and intercrop) were also computed, and selection overlap among top-ranked clones (5%, 10%, 20%) quantified agreement in selection decisions.

### 2.9 Tester comparison

Within each tester, a model equivalent to the intercrop form of Eq. 1 was fitted, with clone, replicate within year, and an AR1 × AR1 residual. Tester mean differences were assessed by Wald F-tests, and the clone × tester interaction by a likelihood-ratio test with a 0.5 boundary correction. Genetic correlations between testers were estimated with bivariate models, and clone BLUPs compared by Pearson and Spearman correlations and Jaccard overlap among top-ranked clones. Broad-sense heritability and expected gain were estimated per tester.

### 2.10 Selection efficiency and gain transfer from monoculture to intercrop

Chance-corrected selection efficiency between monoculture and intercrop followed Hamblin and Zimmermann (1986):

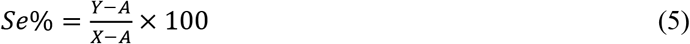

where is the number of clones selected at a given selection intensity, is the number also selected in the alternate system, and is the number of coincidences expected by chance. Selection was evaluated at 5, 10, 15, 20, 25, and 33% intensities using BLUP rankings from the independently fitted monoculture and intercrop models.

To quantify the breeding consequence of indirect selection, gain transfer was calculated by selecting clones in one system and evaluating their predicted performance in the alternate system. Gain transferred from monoculture to intercrop was expressed as a percentage of the gain obtained by direct selection in intercrop, with the reciprocal calculation performed for intercrop-to-monoculture selection. A combined strategy was additionally evaluated by selecting clones on the mean of their monoculture and intercrop BLUPs and expressing the resulting gain in each system relative to direct selection in that system. Results at 10 and 33% selection intensities were summarized for comparison.

### 2.11 Mixing ability, producer and associate effects

General and specific mixing ability were estimated following Haug et al. (2023):

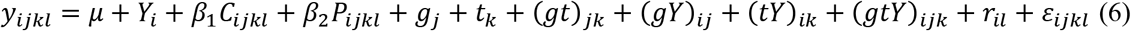

where *g_j_* is the cassava clone effect (general mixing ability), *t_k_* the cowpea tester effect, (*gt*)*_jk_* the specific mixing ability, and (*gY*)*_ij_*, (*tY*)*_ik_*, (*gtY*)*_ijk_* their year interactions; other terms are as in Eq. 1. Thus 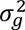 is the general mixing ability variance and 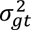 the specific mixing ability variance. The response was the combined intercrop-merit index defined in Section 2.4. Producer and associate effects were obtained from the same model by replacing the response with standardized cassava intercrop yield (producer) or standardized cowpea grain yield (associate).

### 2.12 Cassava partial and total intercrop Land equivalent ratios

For the combined year analysis, cassava partial land equivalent ratio (*pLER*) and total land equivalent ratio (LER) were calculated from spatially adjusted means, comparing each cassava clone’s intercrop yield with its own monoculture yield and each cowpea tester’s intercrop yield with the monoculture yield of the same tester (Crookston and Hill, 1979). Land equivalent ratio was computed from clone means as:

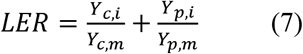

While cassava partial land equivalent ratios were obtained using:

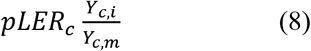

where *Y_c_*_,*i*_and *Y_p_*_,*i*_are cassava and cowpea intercrop yields and *Y_c_*_,*m*_, *Y_p_*_,*m*_the corresponding monoculture yields, respectively.

## 3 Results and discussion

### 3.1 Intercropping reduces cassava yield but maintains a land-use advantage

The two seasons accumulated similar thermal time (year 1, 14,600 °Cd; year 2, 13,800 °Cd) and total rainfall ( approximately 1,750 mm) but differed in rainfall distribution, with only 3 mm during December 2023 to January 2024 and an elevated vapor pressure deficit (up to 1.2 kPa) in year 1 against a shorter low-rainfall window in year 2 (Fig. S1). This contrast provides the environmental differences underlying the genotype-by-year effects reported below.

Pooled across years, monoculture fresh root yielded 30.52 kg plot⁻¹ compared with 24.69 kg plot⁻¹ under intercropping, a reduction of about 5.8 kg plot⁻¹ (19%) (Fig. S2A). The penalty was similar within years, with fresh root yield reduced from 28.98 to 23.22 kg plot⁻¹ in 2023 and from 32.07 to 26.16 kg plot⁻¹ in 2024. This consistent reduction agrees with earlier cassava–legume studies (Cenpukdee and Fukai 1992a; Dapaah et al. 2003; Hidoto and Loha 2013; Nyi et al. 2014; Mwebaze et al. 2024) and indicates a persistent competitive effect of intercropping rather than a response confined to one season. Fresh root yield was consistently lower under intercropping in both years (Fig. S3A).

The cropping-system effect was much smaller for harvest index and negligible for dry matter content (Fig. S2B, C; Fig. S3B, C). Across years, harvest index declined from 0.56 under monoculture to 0.53 under intercropping, whereas dry matter content was essentially unchanged (32.54% vs 32.53%) (Fig. 2B, C). The year-specific patterns were similar, although dry matter content varied substantially between years (Fig. S3B, C). These results suggest that the intercrop penalty primarily affected cassava root yield rather than root dry matter concentration, with only a modest change in biomass partitioning, and highlight the need to breed against cassava’s competitive disadvantage when intercropped (Moore et al. 2022; Annicchiarico et al. 2021).

**Figure 2.**
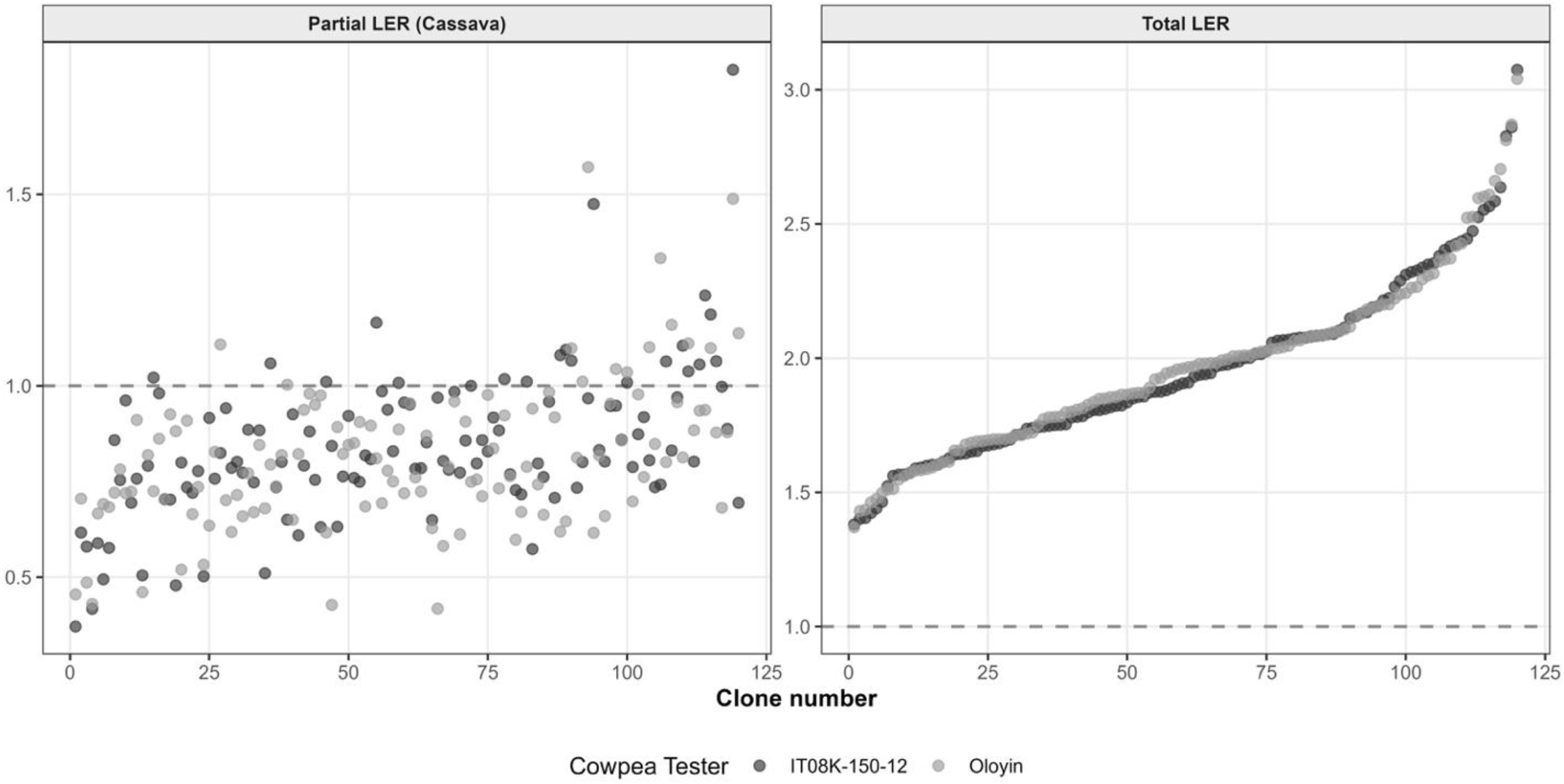
Cassava partial and total land equivalent ratios under cassava–cowpea intercropping. Two scatter-plot panels sharing an x-axis labelled clone number (0 to about 120), with points coloured by cowpea tester: dark grey for IT08K-150-12 and light grey for Oloyin. In both panels clones are sorted left to right by increasing value and a dashed horizontal reference line marks LER equal to 1. Left panel, cassava partial LER: most points fall below the dashed line, rising from about 0.35 at the low end to around 1.0 at the high end, with a scattered subset of clones above 1.0 and a few reaching 1.5 to 1.8; the two testers are intermixed throughout. Right panel, total LER: every point lies well above the dashed line, forming a smooth ascending curve from about 1.4 to just above 3.0, with the two testers overlapping almost completely. The contrast shows cassava alone is usually below break- even while the cassava–cowpea system is always above it.

Within intercropping, cassava fresh root yield differed little between testers, averaging 25.04 kg plot⁻¹ with IT08K-150-12 and 24.34 kg plot⁻¹ with Oloyin across years; similarly small differences occurred within each year (Fig. S2A,B,C; Fig. S3A,B,C). Thus, the cropping-system effect was substantially larger than the effect of cowpea tester identity.

Despite the 19% cassava root-yield penalty, the intercrop maintained a clear land-use advantage. In the combined-year analysis, total land equivalent ratios exceeded 1.0 for all clone–tester combinations, ranging from approximately 1.4 to above 3.0 under both cowpea testers (Fig. 2). This land-use advantage is consistent with broader evidence that intercropping can raise productivity in long-term systems (Yin et al. 2026; Zustovi et al. 2024). Also, the coexistence of reduced cassava yield and greater system-level land-use efficiency is consistent with previous cassava–legume studies and recent meta-analyses, which similarly report cassava yield penalties alongside total LER values above unity (Dapaah et al. 2003; Dettweiler et al. 2023; Aisien et al. 2026). Cassava partial LER was generally below 1.0, consistent with the fresh-root-yield penalty, although a subset of clones approached or exceeded unity (Fig. 2). These genotypes are particularly relevant for breeding because they indicate that some cassava backgrounds can maintain near-monoculture root productivity under intercropping while contributing to a land-efficient cassava–cowpea system.

Although cowpea served primarily as a tester, intercrop grain yield was similar for IT08K-150-12 and Oloyin (255 and 242 kg ha⁻¹, respectively; Fig. S4). Both testers showed lower grain yield in the second season (185 and 161 kg ha⁻¹, respectively) than in the first (288 and 298 kg ha⁻¹), indicating a similar seasonal response across testers. Monoculture means were 231 and 212 kg ha⁻¹, respectively, but these values are presented only as reference because monoculture replication was insufficient for formal statistical comparison. Overall, the comparable performance of the two testers suggests that seasonal conditions exerted a stronger influence on cowpea yield than tester identity, consistent with the limited tester dependence observed for cassava.

### 3.2 Exploitable genetic variation and genotype-by-management interaction

Significant genetic variance among cassava clones was detected for all three traits under both systems across years (Table 1), satisfying the first prerequisite for intercrop breeding (Moore et al. 2022; Haug et al. 2021; Dubey et al. 2024). For fresh root yield, genetic variance was 23.21 under monoculture and 15.49 under intercrop, with moderate broad-sense heritabilities of 0.55 and 0.50 (P < 0.001); year-specific estimates were substantially higher in year 2 (H² = 0.72 monoculture, 0.64 intercrop) than year 1 (0.32 and 0.36). Harvest index showed high heritability across systems (H² = 0.81 monoculture, 0.73 intercrop; P < 0.001) and dry matter content moderate to high heritability (0.68 and 0.75), each with significant clone variance in both systems and years. Intercrop heritabilities are comparable to the mixture-yield heritability of 0.59 reported for pea-barley (Haug et al. 2023).

**Table 1.**
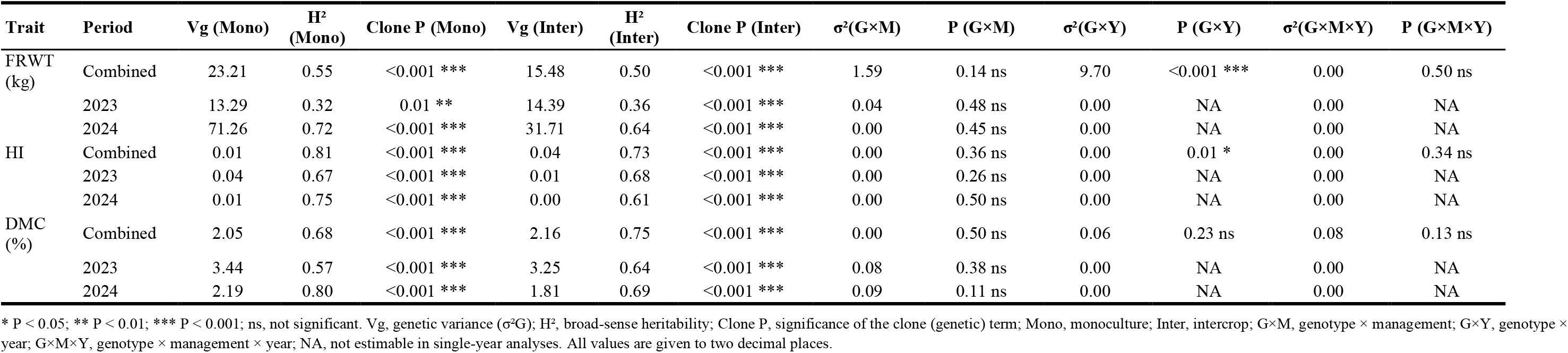
Genetic variance, broad-sense heritability, and genotype-interaction components for three cassava traits under monoculture and intercropping, estimated across years (combined) and within each year. Traits are fresh root weight (FRWT, kg plot⁻¹), harvest index (HI), and root dry matter content (DMC, %). Vg, genetic variance (σ²G); H², entry-mean broad-sense heritability; Clone P, significance of the clone (genetic) term; Mono, monoculture; Inter, intercrop. Interaction variances are genotype × management (σ²G×M), genotype × year (σ²G×Y), and genotype × management × year (σ²G×M×Y), each with its likelihood-ratio test P-value. Significance: *P < 0.05; **P < 0.01; ***P < 0.001; ns, not significant; NA, not estimable in single-year analyses. Values are given to two decimal places.

In the combined model, genotype-by-management (G×M) variance was small and non-significant for all traits, whereas genotype-by-year (G×Y) was significant for fresh root yield (σ²_G×Y_ = 9.69; P < 0.001) and harvest index (σ²_G×Y_ = 0.0004; P = 0.011); no three-way G×M×Y interaction was detected.

The small, non-significant G×M is consistent with monoculture-based selection remaining the primary early-stage strategy, whereas significant G×Y highlights the importance of testing across years and environments. The absence of G×M should not, however, be read as an absence of re-ranking, as the BLUP correlations below demonstrate. Consistently non-significant G×M may reflect the temporal niche separation between cassava (12 months) and cowpea (3 to 4 months), although Zimmermann (1996) found G×M in bean-maize to be environment-dependent, so multi-environment validation is needed before generalizing. These results support the view that minor adaptations to existing monoculture pipelines are a practical first step toward intercrop improvement (Dubey et al. 2024).

### 3.3 Predicting intercrop performance from monoculture: genetic correlation, selection efficiency, and gain transfer

We characterized the monoculture–intercrop relationship using two complementary measures (Table 2): the bivariate genetic correlation (*r_g_*), which estimates the underlying genetic relationship between systems, and Pearson and Spearman correlations between separately fitted monoculture and intercrop clone BLUPs (*r_BLUP_*), which reflect the ranking agreement a breeder would observe. Selection efficiency and predicted gain transfer were then used to quantify the breeding consequences of this agreement.

**Table 2.** Cross-system genetic correlation, selection efficiency, and gain transfer for three cassava traits evaluated under monoculture and cassava–cowpea intercropping, across years (combined) and within each year (2023–2024). Traits are fresh root weight (FRWT, kg plot⁻¹), harvest index (HI), and root dry matter content (DMC, %). H²(M) and H²(I), broad-sense heritability under monoculture and intercrop; r_g, bivariate genetic correlation between systems; r and ρ, Pearson and Spearman correlations between monoculture and intercrop clone BLUPs; Se% (10) and Se% (33), chance-corrected selection efficiency (Hamblin and Zimmermann 1986) at 10% and 33% selection intensities; RG, gain transfer at 10% intensity, expressed as a percentage of the reference direct-selection gain. RG M→I, selecting in monoculture and evaluating in intercrop (relative to direct intercrop selection); RG I→M, the reciprocal; RG Mn→M and RG Mn→I, selecting on the mean of the two systems, relative to direct monoculture and direct intercrop selection, respectively. ***P < 0.001. Values are given to two decimal places.

| Trait | Period | H<sup>2</sup>(M) | H<sup>2</sup>(I) | $r_g$ | $r$ | $\rho$ | Se% (10) | Se% (33) | RG M→I | RG I→M | RG Mn→M | RG Mn→I |
| --- | --- | --- | --- | --- | --- | --- | --- | --- | --- | --- | --- | --- |
| FRWT (kg) | Comb. | 0.59 | 0.59 | 0.99 | 0.652 *** | 0.63 *** | 44.4 | 43.8 | 68.8 | 69.6 | 87.3 | 91.9 |
|  | 2023 | 0.32 | 0.36 | 0.99 | 0.446 *** | 0.42 *** | 25.9 | 21.3 | 31.9 | 60.1 | 74.7 | 97.4 |
|  | 2024 | 0.72 | 0.64 | 0.92 | 0.661 *** | 0.63 *** | 25.9 | 32.5 | 50.5 | 57.1 | 90.6 | 87.1 |
| HI | Comb. | 0.81 | 0.77 | 0.99 | 0.834 *** | 0.85 *** | 72.2 | 66.2 | 89.2 | 87.1 | 97.5 | 93.2 |
|  | 2023 | 0.67 | 0.68 | 0.99 | 0.713 *** | 0.72 *** | 44.4 | 58.8 | 75.6 | 58.0 | 91.1 | 90.4 |
|  | 2024 | 0.75 | 0.61 | 0.99 | 0.771 *** | 0.76 *** | 53.7 | 55.0 | 75.8 | 80.7 | 93.5 | 95.2 |
| DMC (%) | Comb. | 0.70 | 0.76 | 0.99 | 0.845 *** | 0.85 *** | 35.2 | 70.0 | 73.9 | 75.3 | 94.9 | 89.7 |
|  | 2023 | 0.57 | 0.64 | 0.96 | 0.620 *** | 0.61 *** | 25.9 | 43.8 | 68.3 | 53.7 | 91.5 | 89.3 |
|  | 2024 | 0.80 | 0.69 | 0.95 | 0.772 *** | 0.80 *** | 44.3 | 57.9 | 72.6 | 67.8 | 95.2 | 89.3 |

Comb. = combined-year analysis; Pearson = $r$ , Spearman = $\rho$ (BLUP correlations); bivariate $r_g$ ; Hamblin and Zimmermann (1986) Se% at 10% and 33% selection intensities; realized gain (RG%) at 10% intensity. RG M→I = realized gain from selecting in monoculture and testing in intercrop, as % of direct intercrop selection gain; RG I→M = the reverse; RG Mn→M and RG Mn→I = gain from selecting on the system mean, relative to direct monoculture and intercrop selection, respectively. \*\*\* $P < 0.001$ .

Bivariate genetic correlations were high for the focal traits (–0.99; Table 2), including near-unity estimates for fresh root yield and estimates ≥0.95 for harvest index and dry matter content. Robustness analyses across four additional traits: shoot weight, root number, root size, and number of rotten roots, showed a similar pattern: ranged from 0.78 to 1.00, and likelihood-ratio tests failed to reject for any trait- period combination (Table S2). Nevertheless, Spearman agreement between independently estimated monoculture and intercrop BLUPs for these traits ranged from 0.28 to 0.79, and top-10% selection overlap ranged from only 11 to 58%. Thus, the generally high estimates indicate substantial shared genetic signal between systems, but the more modest rank agreement and incomplete elite-set overlap show that high, or even near-unity, should not be interpreted as evidence of identical genotype rankings. Estimates approaching the correlation boundary therefore warrant cautious interpretation.

This distinction was also evident for the focal traits. For fresh root yield, combined-year BLUP agreement was moderate (Pearson 0.65, Spearman 0.63), with only about half of the clones selected at stringent selection intensities shared between systems and rank shifts exceeding 50 positions for some clones (Figure not shown). Harvest index and dry matter content showed stronger BLUP concordance and greater selection overlap. At 10% selection intensity, chance-corrected selection efficiency was 44.4% for fresh root yield in the combined analysis and 25.9% within individual years. More importantly, monoculture-selected clones captured 68.8% of the predicted gain attainable from direct intercrop selection for fresh root yield, with reciprocal gain transfer of similar magnitude (approximately 60–70%). Harvest index transferred more strongly, with 72.2% selection efficiency and 89.2% of direct intercrop gain recovered from monoculture selection, whereas dry matter content was intermediate, with selection efficiency of 25.9–44.3% and gain transfer of approximately 68–74%. Selection on mean performance across the two systems captured 87–92% of direct gain, indicating that combining information from monoculture and intercrop evaluations recovered substantially more of the available selection response than reliance on either system alone.

The gap between the high estimated and more moderate is important because the two quantities describe different aspects of the monoculture–intercrop relationship. The bivariate characterizes the estimated underlying genetic covariance structure, whereas correlations between independently estimated BLUPs additionally reflect finite prediction precision and environmental variation (Lynch and Walsh 1998; Piepho et al. 2008). Estimates approaching the correlation boundary also warrant cautious interpretation because the data may provide limited information for distinguishing very high correlations from unity (Schaeffer 2018). Consequently, particularly when approaches its parameter boundary, rank agreement, elite-set overlap, selection efficiency, and gain transfer provide important complementary evidence of the practical correspondence between systems. Significant genotype-by-year interaction for fresh root yield further indicates temporal re-ranking and helps explain why strong underlying genetic association did not translate into complete selection coincidence across systems, consistent with incomplete selection agreement reported for bean–maize mixtures by Zimmermann (1996).

For breeding practice,, selection efficiency, and gain transfer therefore provide more direct measures of the consequences of using monoculture performance to select for intercropping than alone. The positive cross-system genetic relationship creates favorable conditions for indirect selection (Holland and Brummer 1999; Moore et al. 2022), but the 44.4% selection efficiency and 68.8% gain transfer for fresh root yield show that high does not guarantee complete recovery of superior intercrop genotypes.

Conversely, the 87–92% gain captured by selection on mean performance across systems indicates that strategically combining information from both systems can retain most of the available gain. These results support a staged breeding strategy in which monoculture evaluation remains useful for early selection, followed by targeted intercrop testing of advanced material to recover high-performing clones missed by indirect selection (Haug et al. 2023; Hohmann et al. 2026).

### 3.4 Limited influence of cowpea tester on cassava performance and selection

Cassava performance was broadly consistent across the two testers (Table 3). Tester effects were non-significant for fresh root yield and harvest index in the combined analysis (predicted means 24.72 vs 24.45 kg plot⁻¹ and 0.53 vs 0.52; Table 3), with comparable heritabilities and predicted gains; a small but significant difference appeared only for dry matter content (32.33% vs 32.60%; P = 0.021). Clone-by- tester interaction was non-significant for all traits except fresh root yield in year 2, genetic correlations between testers were high (r_g = 0.995 to 0.998), and top-20% selection overlap was moderate (Jaccard ≈ 0.46 to 0.55; Table S3).

**Table 3.** Effect of cowpea tester identity on cassava performance and selection under intercropping, for three traits across years (combined) and within each year (2023–2024). Traits are fresh root weight (FRWT, kg plot⁻¹), harvest index (HI), and root dry matter content (DMC, %). Mean IT08K and Mean Oloyin, model-adjusted cassava means under the two cowpea testers (IT08K = IT08K-150-12); Tester diff (P), Wald F-test for the tester effect; H², broad-sense heritability under each tester; Exp. gain, expected selection gain (R = i · H² · σ_g) at intensity i = 1.4 (top ∼10%); σ²(G×T) and P(G×T), clone × tester interaction variance and its likelihood-ratio-test significance (0.5 × χ²(1) boundary correction). Predicted means are from mixed models with an AR1 × AR1 spatial residual structure and plant-stand covariates. Significance: *P < 0.05; **P < 0.01; ***P < 0.001; ns, not significant.

| Trait | Period | Mean IT08K | Mean Oloyin | Tester diff (P) | H<sup>2</sup> IT08K | H<sup>2</sup> Oloyin | Exp. gain IT08K | Exp. gain Oloyin | $\sigma^2(G \times T)$ | P (G × T) |
| --- | --- | --- | --- | --- | --- | --- | --- | --- | --- | --- |
| FRWT (kg) | Combined | 24.72 | 24.45 | 0.56 ns | 0.45 | 0.487 | 2.31 | 2.72 | 1.50 | 0.18 ns |
|  | 2023 | 23.72 | 22.73 | 0.15 ns | 0.22 | 0.45 | 0.89 | 2.71 | 0.00 | 0.49 ns |
|  | 2024 | 25.91 | 26.09 | 0.78 ns | 0.63 | 0.74 | 4.69 | 6.78 | 7.92 | 0.01 ** |
| HI | Combined | 0.53 | 0.52 | 0.32 ns | 0.65 | 0.70 | 0.05 | 0.06 | 0.00 | 0.50 ns |
|  | 2023 | 0.51 | 0.50 | 0.45 ns | 0.66 | 0.63 | 0.07 | 0.05 | 0.00 | 0.50 ns |
|  | 2024 | 0.55 | 0.54 | 0.42 ns | 0.64 | 0.58 | 0.06 | 0.05 | 0.04 | 0.10 ns |
| DMC (%) | Combined | 32.33 | 32.60 | 0.02 * | 0.67 | 0.67 | 1.44 | 1.32 | 0.00 | 0.49 ns |
|  | 2023 | 34.76 | 34.81 | 0.78 ns | 0.68 | 0.69 | 1.86 | 1.77 | 0.41 | 0.09 ns |
|  | 2024 | 29.99 | 30.34 | 0.02 * | 0.66 | 0.76 | 1.27 | 1.47 | 0.00 | 0.49 ns |

Cassava means under two cowpea testers (IT08K = IT08K-150-12 and Oloyin); Wald F-tests for tester effects; broad-sense heritability (H<sup>2</sup>); expected selection gain ( $R = i \cdot H^2 \cdot \sigma_g$ ) at selection intensity $i = 1.4$ (top ~10%); and clone × tester interaction for fresh root weight (FRWT, kg plot<sup>-1</sup>), harvest index, and root dry matter content (DMC, %). Results are presented for combined years (2023–2024) and individual years. Clone × tester interaction was tested using likelihood-ratio tests (LRT) with the $0.5 \times \chi^2(1)$ boundary correction appropriate for variance components. Predicted means are model-adjusted estimates from mixed models fitted with AR1 × AR1 spatial residual structure and plant-stand covariates. Significance codes: ns = not significant; \* P < 0.05; \*\* P < 0.01; \*\*\* P < 0.001.

The two contrasting testers thus produced broadly similar overall cassava performance, although agreement among the highest-ranked clones was only moderate. Together with generally non-significant clone-by-tester interaction and high between-tester genetic correlations, these results indicate limited tester dependence in this population, consistent with a GMA-dominant regime and with Wright’s (1985) random-partner assumption. Given only two testers and limited power to detect specific interactions, however, additional testers and environments are needed before concluding that a single tester is sufficient. The result is nonetheless operationally encouraging; standard testers already serve this role in forage and cereal-legume systems, where a small tester set reliably ranks focal-crop genotypes without duplicating the full testing network (Haug et al. 2023; Hohmann et al. 2026), and our findings suggest that a small cowpea tester set could serve the same function in a cassava “advanced-intercrop-yield-trial”.

### 3.5 Yield-scale asymmetry and the determination of intercrop merit

Cassava and cowpea differed markedly in yield magnitude (Fig. 3A): cassava fresh root yield ranged from 5 to 58 kg plot⁻¹ (mean 23.6 kg), whereas cowpea grain yield ranged from 100 to 899 g plot⁻¹ (mean 366.7 g), corresponding to a 64.3-fold difference when expressed on a common mass scale.

**Figure 3.**
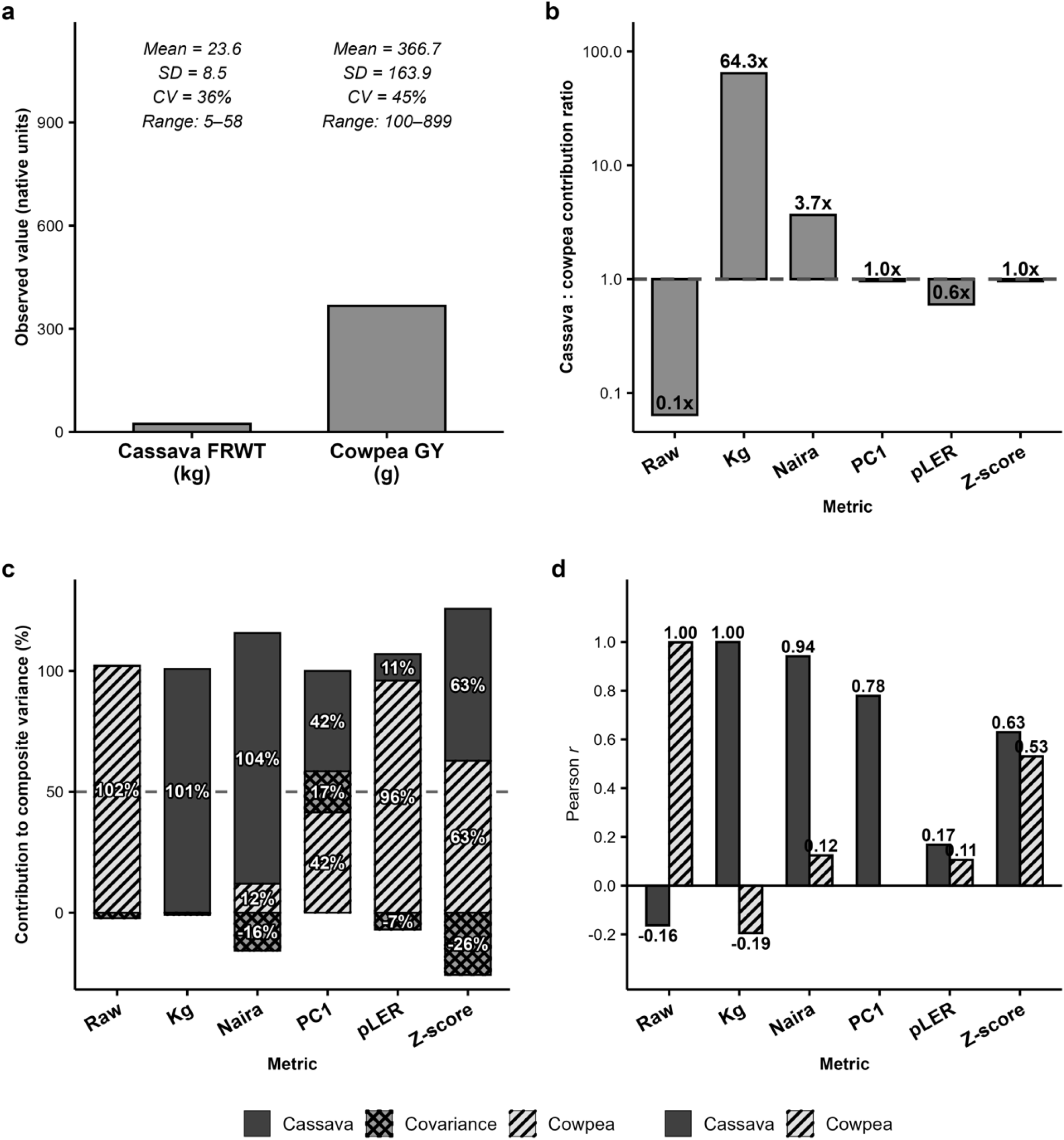
Yield-scale asymmetry between cassava and cowpea and its effect on composite intercrop-merit metrics. Effect of the cassava–cowpea yield-scale asymmetry on six composite intercrop-merit metrics, combined-year data at Ibadan, Nigeria. (a) Observed cassava fresh root weight (FRWT, kg plot⁻¹) and cowpea grain yield (GY, g plot⁻¹) on their native scales, with mean, standard deviation (SD), coefficient of variation (CV) and range; the two crops differ ≈64-fold in numerical magnitude. (b) Cassava-to-cowpea contribution ratio for each metric on a log scale; the dashed line at 1.0 marks equal contribution. (c) Decomposition of composite variance (% of total) into cassava (solid dark), cowpea (hatched) and covariance (cross-hatched) components; the dashed line marks 50%. (d) Pearson correlation of each composite metric with cassava (solid) and cowpea (hatched) component yields. The six metrics are the sum in native units (Raw), the sum in a common mass unit (Kg), an economic-value index (Naira), the first principal component of standardized yields (PC1), a partial-land-equivalent-ratio index (pLER), and a z-score index (Z-score). Native-unit and mass-unit sums were dominated by whichever crop had the larger numerical scale (cowpea and cassava, respectively), whereas the z-score composite balanced the two crops’ contributions and retained positive correlations with both component yields (r = 0.63 and 0.53), justifying its use as the intercrop-merit phenotype

Consequently, the relative contribution of each crop to composite intercrop merit depended strongly on metric construction (Fig. 3B-D). Using yields in their native units favored cowpea (cassava-cowpea contribution ratio = 0.1), with the resulting index almost perfectly correlated with cowpea yield (r = 1.00) but negatively correlated with cassava yield (r = −0.16). Expressing both yields in kilograms reversed this dominance, producing a 64.3-fold cassava bias and an index perfectly correlated with cassava yield (r = 1.00) but negatively correlated with cowpea yield (r = −0.19). Economic weighting reduced, but did not eliminate, the asymmetry (3.7-fold cassava bias; r = 0.94 with cassava and 0.12 with cowpea). In contrast, PC1 and z-score standardization produced equal cassava-cowpea mean contribution ratios, while the z-score index retained substantial positive associations with both cassava and cowpea yields (r = 0.63 and 0.53, respectively). Metric choice also altered estimated genetic parameters and the relative contributions of producer and associate effects (Table S4), as well as genotype rankings . These results justified use of the z-score composite for subsequent mixing-ability analyses because it prevented either crop from dominating intercrop merit solely because of differences in measurement scale.

Mathematically, summing standardized component yields is equivalent, apart from an additive constant, to weighting each component inversely by its standard deviation. This scale adjustment is consistent with the broader use of weighted selection indices to regulate the relative contributions of component species (Sampoux et al. 2020), while retaining the separated-yield perspective used in mixture-breeding frameworks such as Haug et al. (2023). Explicit standardization was particularly important in the cassava–cowpea system because the large disparity in component yield scales caused unstandardized metrics to be dominated by whichever crop had the larger numerical scale. Thus, standardization provided a more balanced definition of intercrop merit for evaluating the genetic contributions of cassava producers and cowpea associates in the subsequent mixing-ability analysis.

### 3.6 Producer-dominated mixing ability

Clone GMA was significant for the trait-specific merit index across all three traits (fresh root yield LRT = 7.83, P = 0.003; harvest index LRT = 26.21, P < 0.001; dry matter content LRT = 26.02, P < 0.001; Table 4), while SMA was negligible in the combined-year and 2023 analyses, reaching significance only for harvest index and dry matter content in 2024 (LRT = 3.05 and 4.43, both P < 0.05). The cassava producer effect model (using the cassava trait as the response variable) produced even larger clone effects (fresh root yield LRT = 16.12; harvest index 55.06; dry matter content 74.59; all P < 0.001).

**Table 4.** Mixing-ability parameters for the cassava–cowpea intercrop analysis across the combined-year dataset and each individual year. GMA was estimated on the trait-specific merit index (focal cassava trait z-score + cowpea grain z-score); Producer on the focal cassava trait; Associate on cowpea grain yield. LRT and significance are from likelihood-ratio tests of each random effect against a reduced model, referred to a 0.5:0.5 mixture of χ²(0) and χ²(1). H² = entry-mean broad-sense heritability; clone variance (%) = clone share of total phenotypic variance. Associate is a single cowpea-grain analysis, so its values are common across cassava traits within a year, differing for dry matter content only where plots lacking a dry matter record are absent. ns, not significant; * P < 0.05; ** P < 0.01; *** P < 0.001.

| Parameter | Combined | 2023 | 2024 |
| --- | --- | --- | --- |
| **Fresh root yield** |  |  |  |
| GMA (Clone) LRT | 7.83** | 2.95* | 10.13*** |
| GMA $H^2$ | 0.34 | 0.15 | 0.48 |
| GMA clone variance (%) | 11.84 | 7.96 | 27.49 |
| SMA (Clone × Cowpea) LRT | 0.00 ns | 0.00 ns | 0.10 ns |
| Producer (Clone) LRT | 16.12*** | 15.55*** | 13.47*** |
| Producer $H^2$ | 0.47 | 0.36 | 0.63 |
| Producer clone variance (%) | 20.35 | 21.48 | 37.97 |
| Associate (Clone) LRT | 0.14 ns | 5.81** | 0.39 ns |
| Associate $H^2$ | 0.05 | 0.20 | 0.28 |
| Associate clone variance (%) | 1.36 | 10.89 | 4.57 |
| **Harvest index** |  |  |  |
| GMA (Clone) LRT | 26.21*** | 43.69*** | 7.81** |
| GMA $H^2$ | 0.55 | 0.53 | 0.56 |
| GMA clone variance (%) | 26.20 | 35.89 | 24.46 |
| SMA (Clone × Cowpea) LRT | 0.00 ns | 0.00 ns | 3.05* |
| Producer (Clone) LRT | 55.06*** | 80.25*** | 10.65*** |
| Producer $H^2$ | 0.72 | 0.70 | 0.60 |
| Producer clone variance (%) | 41.90 | 53.49 | 36.86 |
| Associate (Clone) LRT | 0.14 ns | 5.81** | 0.39 ns |
| Associate $H^2$ | 0.05 | 0.20 | 0.28 |
| Associate clone variance (%) | 1.36 | 10.89 | 4.57 |
| **Dry matter content** |  |  |  |
| GMA (Clone) LRT | 26.02*** | 30.57*** | 14.36*** |
| GMA $H^2$ | 0.56 | 0.52 | 0.67 |
| GMA clone variance (%) | 26.86 | 32.34 | 33.24 |
| SMA (Clone × Cowpea) LRT | 0.05 ns | 0.24 ns | 4.43* |
| Producer (Clone) LRT | 74.59*** | 47.92*** | 28.45*** |
| Producer $H^2$ | 0.75 | 0.69 | 0.80 |
| Producer clone variance (%) | 46.72 | 44.09 | 54.30 |
| Associate (Clone) LRT | 0.11 ns | 5.81** | 0.09 ns |
| Associate $H^2$ | 0.04 | 0.20 | 0.29 |
| Associate clone variance (%) | 1.23 | 10.89 | 2.33 |

GMA, general mixing ability; SMA, specific mixing ability; LRT, likelihood-ratio test statistic; $H^2$ , entry-mean broad-sense heritability; clone variance (%), clone share of total phenotypic variance; ns, not significant. Significance of each random effect from likelihood-ratio tests referred to a 0.5:0.5 mixture of $\chi^2(0)$ and $\chi^2(1)$ : \* $P < 0.05$ ; \*\* $P < 0.01$ ; \*\*\* $P < 0.001$ .

Heritability followed the same order (Table 4): producer effects 0.47 to 0.75, GMA 0.34 to 0.56, and associate effects consistently low (0.04 to 0.05). Variance decomposition (Fig. S5) showed clone effects explaining 11.8 to 26.9% of total variance under the GMA model but 20.4% (fresh root yield), 41.9% (harvest index) and 46.7% (dry matter content) under the producer model, against only 1.2 to 1.4% under the associate model; genetic signals strengthened in year 2, where GMA heritability rose to 0.48 to 0.67.

Predominantly negligible SMA with producer-dominated GMA indicates predominantly general rather than partner-specific mixing ability, a condition expected to reduce the number of testers required for effective evaluation (Annicchiarico et al. 2019; Moore et al. 2022; Hohmann et al. 2026); because the number of possible cassava-by-cowpea combinations is too large for exhaustive evaluation, negligible SMA converts an otherwise intractable factorial into a manageable advanced-stage screen. The same producer-dominated, SMA-absent structure was reported for pea-barley (Haug et al. 2023) and maps onto the Sampoux et al. (2020) scenario in which parallel GMA selection is as efficient as reciprocal selection, though the two-tester design has limited power for SMA detection (Haug et al. 2021), so this conclusion is drawn with appropriate caution.

Plotting producer against associate BLUPs (Fig. 4) following Haug et al. (2021) distributed clones across all four quadrants, with 22 to 27% in the mutualistic Pr+ As+ sector that Wright (1985) and Haug et al. 2021 identified as optimal for economic yield, indicating that favorable allele combinations already segregate at appreciable frequency (Haug et al. 2023). The producer-to-associate effects trade-off was weaker than in pea-barley (r = −0.31 vs −0.65; Haug et al. 2023), therefore selecting for high producer effects should not severely affect associate effects. GMA correlated with producer (r = 0.79), associate (0.52) and monoculture performance (0.57), and producer with monoculture (0.68); for harvest index and dry matter content the producer-to-monoculture correlations were strong (0.85 and 0.82; Fig. S6). Strong producer-to-monoculture coupling validates trait-informed monoculture selection (Moore et al. 2022, Haug et al. 2023), whereas the absence of an associate-to-monoculture correlation confirms that monoculture selection captures only the direct component and that associate effects require direct intercrop evaluation (Barot et al. 2017; Sampoux et al. 2020).

**Figure 4.**
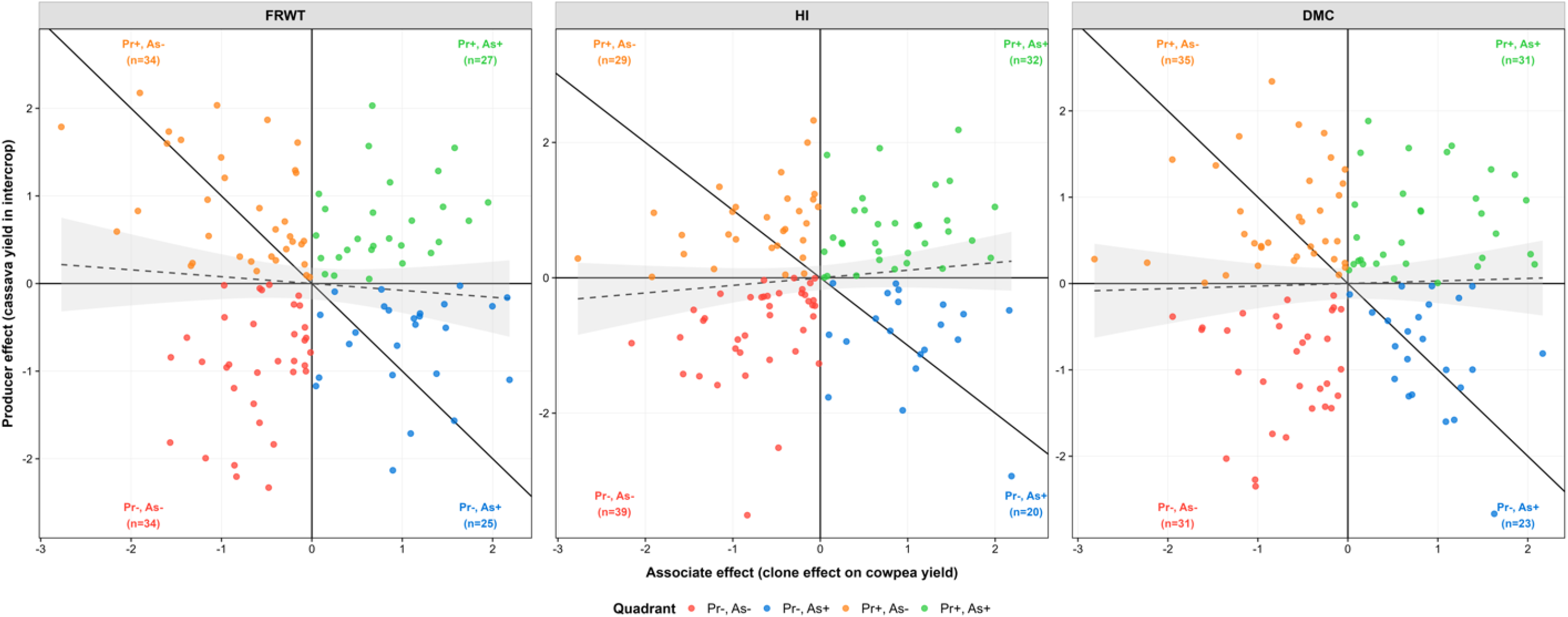
Producer and associate effect BLUPs of cassava clones under cowpea intercropping. Producer effect (a clone’s own cassava yield in intercrop, y-axis) plotted against associate effect (a clone’s effect on cowpea yield, x-axis) for three cassava traits: fresh root weight (FRWT), harvest index (HI), and root dry matter content (DMC), combined-year data at Ibadan, Nigeria. Both effects are clone BLUPs in standardized units. Solid horizontal and vertical lines at zero divide each panel into four quadrants defined by the sign of the two effects, and points are coloured by quadrant: Pr+ As+ (green, favourable for both crops), Pr+ As− (orange), Pr− As+ (blue), and Pr− As− (red, unfavourable for both); the per-quadrant clone counts (n) are printed in each corner. The solid diagonal is the line of equal and opposite effects, and the dashed line with its grey 95% confidence band is the producer–associate regression. Clones were distributed across all four quadrants, with 22–27% falling in the mutualistic Pr+ As+ sector, and the producer– associate relationship was weak (near-flat dashed line), indicating that a clone’s ability to maintain its own yield was largely independent of its effect on the companion crop.

In summary, producer-dominated GMA, negligible SMA, and the positive association between producer effects and monoculture performance indicate that cassava intercrop performance in this population depended more strongly on a clone’s ability to maintain its own performance under competition than on partner-specific interactions. This pattern is consistent with mixture-breeding frameworks in which strong producer and general mixing effects favor the use of monoculture information, while limited SMA reduces the need for partner-specific selection (Sampoux et al. 2020; Haug et al. 2023). The incomplete correspondence between systems nevertheless supports targeted intercrop testing of advanced material (Dubey et al. 2024; Hohmann et al. 2026).

### 3.7 Breeding implications: targeted intercrop evaluation within the existing pipeline

The simulation study by Dubey et al. (2024) showed that the monocrop-intercrop genetic correlation, more than heritability, governs the optimal stage for introducing intercrop testing: at a correlation of 0.9 testing can be delayed to advanced or elite yield trials, whereas at 0.3 it should begin at preliminary trials. Although the cross-system genetic correlations were imprecisely resolved near the parameter boundary, their high estimates, together with non-significant G×M and the substantial gain transferred from monoculture selection, are consistent with the high-correlation, late-testing scenario described by Dubey et al. (2024). This strategy also provides a quantitative-genetic basis for the earlier phenotypic observations of Cenpukdee and Fukai (1992b), who proposed initial sole-crop selection of cassava while recognizing that selection requirements depend on the competitiveness of the companion crop. Our results showed that cassava clones from IITA NextGen advanced yield trials, selected under monoculture, still expressed significant intercrop variation, indicating that monoculture selection did not completely remove intercrop-relevant variation. The moderate Se% for fresh root yield (25.9 to 44.4%) nonetheless calls for caution: a confirmatory single-tester, pooled-tester (Annicchiarico et al. 2021) or incomplete-factorial (Haug et al. 2021) intercrop evaluation at the advanced-yield-trial stage would sharpen root-yield selection.

These findings translate recent calls for interaction-based, community-level breeding into an operational strategy for cassava (Haug and Bourke 2025; Hohmann et al. 2026). Recognizing mixing ability as a sustainability-relevant varietal property, breeding for diversified systems should move from isolated-genotype evaluation toward mixture testing while using standard testers to keep experimental complexity manageable (Hohmann et al. 2026). Framed as frugal innovation, this adapts the existing pipeline to a diversified production system rather than building a parallel one (Haug and Bourke 2025). The monoculture pipeline can therefore be preserved and complemented by targeted cowpea-intercrop evaluation at the advanced-yield-trial stage, a minimal, low-cost adaptation aligned with breeding cassava for the land-efficient, drought-resilient cassava-cowpea systems on which African smallholders depend (Adam et al. 2025).

## 4 Conclusion

Here we show, for the first time, that improved performance in a tropical root–legume intercropping system is largely captured by existing monoculture selection, indicating that cassava can be bred for intercropping with minimal additions to current pipelines rather than through a separate breeding program. Intercrop performance was governed primarily by general mixing ability and producer effects, whereas specific mixing ability and clone-by-tester interaction were limited. Producer effects tracked monoculture performance, supporting monoculture as an efficient early-stage selection environment. However, monoculture selection recovered only about 69% of the predicted direct intercrop gain for fresh root yield, indicating that exclusive monoculture selection can forfeit roughly one-third of achievable gain. A staged strategy is therefore indicated: retain monoculture selection in early breeding stages, and introduce targeted intercrop evaluation of advanced material using a limited set of testers. More broadly, these results show how breeding for diversified cropping systems can be incorporated through focused modifications to existing pipelines rather than through experimentally intractable or technologically intensive schemes (Haug and Bourke 2025).

These conclusions are based on one location, two years, and two cowpea testers, limiting inference about genotype × environment × management interactions and the extent of specific mixing ability (Haug et al. 2021). The high between-system genetic correlations were also estimated near the parameter boundary; consequently, inference about the usefulness of monoculture selection rests more strongly on observed BLUP agreement, selection efficiency, and predicted gain transfer than on alone. Validation across additional environments, management conditions, planting densities, and more diverse cowpea testers is therefore needed (Cenpukdee and Fukai 1992c; Brooker et al. 2015; Hohmann et al. 2026). Incomplete-factorial testing and genomic prediction of general mixing ability could further reduce the number of cassava–cowpea combinations requiring field evaluation (Bančič et al. 2021; Wolfe et al. 2021).

## Acknowledgments

The authors thank all staff and interns of the Cassava Breeding Unit at the International Institute of Tropical Agriculture (IITA), Ibadan, Nigeria, for their support during the field experiments.

## Declarations Funding

This study was jointly funded by the Cassava Breeding Program at the International Institute of Tropical Agriculture (IITA), Ibadan, Nigeria, and the Wolfe Lab, Department of Crop, Soil and Environmental Sciences, Auburn University, Alabama, USA.

## Conflicts of interest

The authors declare that they have no known competing financial or non-financial interests or personal relationships that could have appeared to influence the work reported in this article.

## Ethics approval

Not applicable.

## Consent to participate

Not applicable.

## Consent for publication

Not applicable.

## Availability of data and material

The datasets generated and/or analyzed during the current study are available from the corresponding author on reasonable request.

## Code availability

The R code used for data analysis in the current study is available from the corresponding author on reasonable request.

## Authors’ contributions

Conceptualization: O.G.O., I.Y.R., M.W.; Data curation: O.G.O., U.E.; Formal analysis: O.G.O.; Funding acquisition: I.Y.R., M.W.; Investigation: A.T., I.P., O.G.O.; Methodology: O.G.O., O.O.T., A.T., I.P., I.Y.R., M.W.; Project administration: O.G.O., A.T., I.P., I.Y.R., M.W.; Resources: I.Y.R., M.W.; Software: M.W.; Supervision: O.G.O., I.Y.R., M.W.; Validation: I.Y.R., M.W.; Visualization: O.G.O.; Writing original draft: O.G.O.; Writing, review and editing: O.G.O., A.T., I.P., U.E., O.O.T., I.Y.R., M.W.

## AI use statement

Claude AI was used to assist with writing and debugging R code, and Grammarly was used for grammar and language polishing. All final scientific content, analyses, interpretations, and manuscript text were reviewed and validated by the authors.

## Notes

### Competing Interest Statement

The authors have declared no competing interest.

