## Supplementary tables and figures for "Breeding cassava for intercropping with cowpea: monoculture selection captures most intercrop selection gain, but targeted testing remains necessary"

**Table S1.** Comparison of combined-year bivariate models for the genetic correlation between monoculture (M) and intercrop (I) systems, contrasting the genotype-by-year (G×Y) covariance structure and the spatial-grid specification.

Model A: diagonal G×Y with separate per-management grids; B: unstructured G×Y with separate grids; C: diagonal G×Y with a common physical grid; D: unstructured G×Y with a common grid. All models retain an unstructured us(trait):Clone genetic effect and identical fixed effects, so AIC and BIC are comparable across models. A vs B and C vs D are nested (1 df, covariance = 0 at an interior point); A vs C and B vs D are non-nested and compared by AIC/BIC only. Under a diagonal G×Y the across-management covariance is fixed at zero by construction; a dash denotes a parameter not estimated (or a non-converged model, Conv. = no), not an estimate of zero. Model D for fresh root weight approached the correlation boundary. A common grid with a diagonal G×Y structure was retained as the primary model. k, number of free (co)variance parameters; logLik, residual log-likelihood; AIC and BIC, information criteria (lower is better); r_g, estimated genetic correlation between systems; σ² (M) and σ² (I), genotype-by-year variances under monoculture and intercrop; cov, across-management G×Y covariance; r_g,yr (SE), correlation of year-specific genetic deviations across managements with its standard error (estimated only under unstructured G×Y). Separable AR1 × AR1 residual autocorrelations (not shown) ranged 0.01–0.42 along rows and 0.01–0.57 along columns across models. us, unstructured; diag, diagonal.

| Trait | Model | G×Y | Grid | Conv. | k | logLik | AIC | BIC | $r_g$ | $\sigma^2$ (M) | $\sigma^2$ (I) | cov | $r_{g,yr}$ (SE) |
|---|---|---|---|---|---|---|---|---|---|---|---|---|---|
| FRWT (kg) | A | diag | separate | yes | 18 | −3615.29 | 7266.6 | 7361.2 | 0.996 | 6.785 | 7.423 | — | — |
| | B | us | separate | no | — | — | — | — | — | — | — | — | — |
| | C | diag | common | yes | 12 | −3623.63 | 7271.3 | 7334.3 | 0.997 | 15.334 | 6.618 | — | — |
| | D | us | common | yes | 13 | −3609.98 | 7246.0 | 7314.3 | 0.867 | 19.339 | 8.306 | 12.425 | 0.980 (0.177) |
| HI | A | diag | separate | yes | 18 | 2889.97 | −5743.9 | −5649.3 | 0.998 | 0.0002 | 0.0005 | — | — |
| | B | us | separate | no | — | — | — | — | — | — | — | — | — |
| | C | diag | common | yes | 12 | 2892.54 | −5761.1 | −5698.0 | 0.998 | 0.0005 | 0.0005 | — | — |
| | D | us | common | yes | 13 | 2897.23 | −5768.5 | −5700.1 | 0.999 | 0.0007 | 0.0006 | 0.0005 | 0.822 (0.335) |
| DMC (%) | A | diag | separate | yes | 18 | −1503.83 | 3043.7 | 3138.0 | 0.999 | 0.244 | 0.038 | — | — |
| | B | us | separate | yes | 19 | −1503.39 | 3044.8 | 3144.3 | 0.998 | 0.317 | 0.068 | 0.080 | 0.549 (0.606) |
| | C | diag | common | yes | 12 | −1505.90 | 3035.8 | 3098.7 | 0.999 | 0.099 | 0.017 | — | — |
| | D | us | common | no | — | — | — | — | — | — | — | — | — |

Table S2. Genetic correlation, rank agreement, and top-10% selection overlap between monoculture and intercrop performance for four robustness traits across three analysis periods.

Table S2. Between-system relationships for four additional cassava traits across the combined-year, 2023, and 2024 analyses; the three focal traits are reported in Table 2. shwt, shoot weight (kg); rtno, number of roots; rtsz, root size; rtrot, number of rotten roots. *r*g, genetic correlation from the bivariate model; LRT $\chi^2$ (P), likelihood-ratio test of the hypothesis *r*g = 1, referred to a 0.5:0.5 mixture of $\chi^2(0)$ and $\chi^2(1)$; *r*, Pearson and ρ, Spearman correlation between monoculture and intercrop clone BLUPs; top-10% overlap, percentage of clones common to the top 10% selected in each system. ‡ negative LRT statistic: at convergence the constrained model (*r*g = 1) marginally outperformed the unconstrained model and was treated as LRT = 0. Near-unity *r*g estimates lie close to the correlation boundary, where the test has limited power to distinguish *r*g from 1.

| Trait | Period | $r_g$ | LRT $\chi^2$ (P) | $r$ | ρ | Top-10% overlap |
|---|---|---|---|---|---|---|
| shwt | Combined | 1.00 | −0.003‡ (1.000) | 0.763 | 0.792 | 17% |
| | 2023 | 0.78 | 0.000 (0.988) | 0.537 | 0.599 | 17% |
| | 2024 | 0.96 | 0.000 (0.993) | 0.706 | 0.702 | 42% |
| rtno | Combined | 1.00 | 0.003 (0.954) | 0.653 | 0.654 | 42% |
| | 2023 | 0.90 | 0.001 (0.977) | 0.519 | 0.528 | 33% |
| | 2024 | 0.85 | 0.000 (1.000) | 0.581 | 0.556 | 17% |
| rtsz | Combined | 0.88 | 0.000 (0.990) | 0.484 | 0.506 | 33% |
| | 2023 | 1.00 | 0.000 (0.983) | 0.413 | 0.311 | 25% |
| | 2024 | 0.96 | 0.000 (1.000) | 0.464 | 0.438 | 25% |
| rtrot | Combined | 1.00 | 0.002 (0.961) | 0.602 | 0.526 | 58% |
| | 2023 | 1.00 | 0.000 (0.983) | 0.312 | 0.282 | 11% |
| | 2024 | 0.80 | 0.000 (1.000) | 0.373 | 0.393 | 27% |

Shwt, shoot weight (kg); rtno, number of roots; rtsz, root size; rtrot, number of rotten roots. $r_g$, genetic correlation from the bivariate model; r, Pearson and ρ, Spearman correlation between monoculture and intercrop clone BLUPs; top-10% overlap, percentage of clones common to the top 10% selected in each system. ‡ Negative LRT statistic: the constrained model ($r_g$ = 1) marginally outperformed the unconstrained model at convergence and was treated as LRT = 0. Near-unity $r_g$ estimates lie close to the correlation boundary, where the likelihood-ratio test has limited power to distinguish $r_g$ from 1.

Table S3. Tester performance and validation metrics for cassava evaluation under two cowpea testers.

Table S3. Tester performance and validation metrics for cassava evaluation under two cowpea testers (IT08K-150-12 and Oloyin), for fresh root weight (FRWT), harvest index (HI), and root dry matter content (DMC), combined analysis (2023–2024). Mean, predicted mean; $\sigma^2g$, genetic variance; CVg, coefficient of genetic variation (%); Exp. gain, expected selection gain at 10% selection intensity; *r*g(mn), genetic correlation between monoculture and intercrop performance; *r*g(yr), year-to-year genetic correlation (2023 vs 2024); *r*g(testers), bivariate genetic correlation between testers; Jaccard (20%), selection overlap between testers as the Jaccard similarity of the top 20% selected clones. Together these statistics assess tester equivalence, selection stability, and the reliability of intercrop evaluation for cassava breeding.

| Trait | Tester | Mean | $\sigma^2$g | $CV_g$ (%) | Exp. gain | $r_g$(mn) | $r_g$(yr) | $r_g$(testers) | Jaccard (20%) |
|---|---|---|---|---|---|---|---|---|---|
| FRWT | IT08K-150-12 | 24.72 | 13.63 | 14.7 | 2.30 | 0.93 | 0.74 | 0.99 | 0.45 |
| | Oloyin | 24.45 | 16.68 | 16.8 | 2.72 | 0.97 | 0.57 | 0.99 | 0.45 |
| HI | IT08K-150-12 | 0.53 | 0.00 | 11.2 | 0.05 | 0.99 | 0.76 | 0.99 | 0.54 |
| | Oloyin | 0.52 | 0.00 | 11.7 | 0.06 | 0.99 | 0.89 | 0.99 | 0.54 |
| DMC | IT08K-150-12 | 32.33 | 2.34 | 4.7 | 1.44 | 1.00 | 0.91 | 0.99 | 0.45 |
| | Oloyin | 32.60 | 1.98 | 4.3 | 1.31 | 0.99 | 0.89 | 0.99 | 0.45 |

Summary statistics under two cowpea testers (IT08K-150-12 and Oloyin) for fresh root weight (FRWT), harvest index (HI), and root dry matter content (DMC), combined analysis (2023–2024). Mn = monoculture. Mean, predicted mean; $\sigma^2g$, genetic variance; CVg, coefficient of genetic variation (%); Exp. gain, expected selection gain at 10% selection intensity. rg(mn), genetic correlation between monoculture and intercrop performance; rg(yr), year-to-year genetic correlation (2023 vs 2024); rg(testers), bivariate genetic correlation between testers; Jaccard (20%), selection overlap between testers as the Jaccard similarity of the top 20% selected clones. These statistics assess tester equivalence, selection stability, and the reliability of intercrop evaluation for cassava breeding.

Table S4. Model comparison across six intercrop-merit metrics for three focal cassava traits and three analysis periods.

Table S4. Model comparison across six candidate intercrop-merit metrics for three focal cassava traits (fresh root weight, FRWT; harvest index, HI; root dry matter content, DMC) and three analysis periods (combined-year, 2023, 2024). Metrics: Kg, sum of component yields on a common mass scale; Naira, economic-value index from crop-specific prices; PC1, first principal component of standardized component yields; PLER, partial-land-equivalent-ratio index; Raw, sum of yields in native units; Zscore, sum of within-year z-scored component yields. AIC, Akaike information criterion; BIC, Bayesian information criterion (both lower is better); ΔAIC and ΔBIC are relative to the best-fitting metric within each trait-period combination (best = 0). *$H^2$*, broad-sense heritability; GMA%, SMA%, and Res%, percentage of total variance attributable to general mixing ability, specific mixing ability, and residual, respectively. Metric choice for the mixing-ability analyses was based on scale-balance between the two component crops (Section 3.5), not on information criteria alone; the z-score composite was retained because it prevented either crop from dominating intercrop merit through differences in measurement scale.

| Focal trait | Period | Metric | AIC | BIC | ΔAIC | ΔBIC | $H^2$ | GMA% | SMA% | Res% |
|---|---|---|---|---|---|---|---|---|---|---|
| DMC | Combined | Kg | 1714.0 | 1773.4 | 2129.6 | 2129.5 | 0.707 | 42.5 | 0.0 | 44.4 |
| | | Naira | 8583.8 | 8643.2 | 8999.4 | 8999.3 | 0.547 | 21.0 | 2.7 | 56.9 |
| | | PC1 | 233.6 | 293.0 | 649.2 | 649.1 | 0.481 | 19.5 | 2.5 | 61.2 |
| | | PLER | -415.6 | -356.1 | 0.0 | 0.0 | 0.146 | 1.7 | 0.0 | 76.8 |
| | | Raw | 7674.6 | 7734.1 | 8090.2 | 8090.2 | 0.014 | 0.4 | 0.0 | 82.5 |
| | | Zscore | 1065.9 | 1125.3 | 1481.5 | 1481.4 | 0.548 | 25.2 | 1.6 | 58.0 |
| | 2023 | Kg | 1253.7 | 1282.6 | 1902.3 | 1902.4 | 0.812 | 44.1 | 7.8 | 48.1 |
| | | Naira | 5592.8 | 5621.7 | 6241.4 | 6241.5 | 0.718 | 29.7 | 4.7 | 61.1 |
| | | PC1 | 414.6 | 443.5 | 1063.2 | 1063.3 | 0.655 | 25.9 | 6.4 | 67.8 |
| | | PLER | -648.6 | -619.8 | 0.0 | 0.0 | 0.392 | 12.5 | 0.0 | 86.1 |
| | | Raw | 4976.5 | 5005.3 | 5625.1 | 5625.1 | 0.306 | 9.9 | 0.0 | 90.1 |
| | | Zscore | 703.3 | 732.1 | 1351.9 | 1351.9 | 0.675 | 31.4 | 2.8 | 65.8 |
| | 2024 | Kg | 534.6 | 559.4 | 294.2 | 294.2 | 0.897 | 52.9 | 13.7 | 31.6 |
| | | Naira | 2996.8 | 3021.6 | 2756.4 | 2756.4 | 0.767 | 30.6 | 12.3 | 54.9 |
| | | PC1 | 242.5 | 267.3 | 2.1 | 2.1 | 0.752 | 30.3 | 9.9 | 56.9 |
| | | PLER | 240.4 | 265.2 | 0.0 | 0.0 | 0.769 | 3.1 | 17.6 | 54.5 |
| | | Raw | 2668.8 | 2693.7 | 2428.4 | 2428.5 | 0.451 | 2.5 | 14.6 | 82.9 |
| | | Zscore | 392.3 | 417.2 | 151.9 | 152.0 | 0.802 | 33.2 | 17.1 | 49.6 |
| FRWT | Combined | Kg | 3655.6 | 3715.1 | 3754.6 | 3754.5 | 0.470 | 20.3 | 0.0 | 67.7 |
| | | Naira | 10009.9 | 10069.4 | 10108.9 | 10108.8 | 0.483 | 20.9 | 0.0 | 69.0 |
| | | PC1 | 560.9 | 620.5 | 659.9 | 659.9 | 0.298 | 10.7 | 0.4 | 73.5 |
| | | PLER | -99.0 | -39.4 | 0.0 | 0.0 | 0.148 | 0.0 | 0.0 | 75.3 |
| | | Raw | 7728.9 | 7788.4 | 7827.9 | 7827.8 | 0.039 | 1.1 | 0.0 | 82.5 |
| | | Zscore | 1039.4 | 1099.0 | 1138.4 | 1138.4 | 0.341 | 11.8 | 0.0 | 81.7 |
| | 2023 | Kg | 2342.7 | 2371.5 | 2694.0 | 2694.0 | 0.523 | 21.2 | 0.0 | 78.5 |
| | | Naira | 6320.8 | 6349.6 | 6672.1 | 6672.1 | 0.444 | 16.7 | 0.0 | 83.3 |
| | | PC1 | 469.0 | 497.8 | 820.3 | 820.3 | 0.539 | 22.6 | 0.0 | 77.4 |

| Focal trait | Period | Metric | AIC | BIC | ΔAIC | ΔBIC | H² | GMA% | SMA% | Res% |
|---|---|---|---|---|---|---|---|---|---|---|
| | | PLER | -351.3 | -322.5 | 0.0 | 0.0 | 0.456 | 15.1 | 0.0 | 82.7 |
| | | Raw | 4923.7 | 4952.5 | 5275.0 | 5275.0 | 0.308 | 10.0 | 0.0 | 90.0 |
| | | Zscore | 654.0 | 682.8 | 1005.3 | 1005.3 | 0.258 | 8.0 | 0.0 | 92.0 |
| | | Kg | 1333.3 | 1358.5 | 1107.8 | 1107.9 | 0.778 | 38.3 | 8.4 | 53.3 |
| | | Naira | 3687.7 | 3712.8 | 3462.2 | 3462.2 | 0.791 | 41.5 | 7.1 | 51.5 |
| | 2024 | PC1 | 225.5 | 250.6 | 0.0 | 0.0 | 0.577 | 8.7 | 16.7 | 74.5 |
| | | PLER | 261.5 | 286.6 | 36.0 | 36.0 | 0.754 | 6.5 | 11.8 | 56.7 |
| | | Raw | 2775.2 | 2800.3 | 2549.7 | 2549.7 | 0.446 | 6.1 | 10.3 | 83.3 |
| | | Zscore | 390.8 | 415.9 | 165.3 | 165.3 | 0.645 | 27.5 | 2.8 | 68.7 |
| | | Kg | -1977.4 | -1917.8 | 0.0 | 0.0 | 0.427 | 18.1 | 0.0 | 65.6 |
| | | Naira | 8242.2 | 8301.8 | 10219.6 | 10219.6 | 0.134 | 1.1 | 0.0 | 77.0 |
| | Combined | PC1 | 492.4 | 552.0 | 2469.8 | 2469.8 | 0.495 | 20.3 | 1.4 | 68.5 |
| | | PLER | -338.3 | -278.8 | 1639.1 | 1639.0 | 0.168 | 1.9 | 0.0 | 76.0 |
| | | Raw | 7744.2 | 7803.8 | 9721.6 | 9721.6 | 0.038 | 1.1 | 0.0 | 82.2 |
| | | Zscore | 1077.2 | 1136.8 | 3054.6 | 3054.6 | 0.544 | 25.6 | 0.0 | 61.8 |
| | | Kg | -1193.2 | -1164.3 | 0.0 | 0.0 | 0.581 | 25.7 | 0.0 | 74.3 |
| | | Naira | 5250.9 | 5279.8 | 6444.1 | 6444.1 | 0.477 | 10.4 | 0.2 | 81.4 |
| | | PC1 | 412.1 | 440.9 | 1605.3 | 1605.2 | 0.649 | 31.6 | 0.0 | 68.4 |
| HI | 2023 | PLER | -580.9 | -552.1 | 612.3 | 612.2 | 0.383 | 11.6 | 0.0 | 86.5 |
| | | Raw | 4940.1 | 4968.9 | 6133.3 | 6133.2 | 0.328 | 10.9 | 0.0 | 89.1 |
| | | Zscore | 669.7 | 698.5 | 1862.9 | 1862.8 | 0.680 | 34.7 | 0.0 | 65.3 |
| | | Kg | -769.3 | -744.2 | 0.0 | 0.0 | 0.689 | 20.7 | 13.1 | 64.4 |
| | | Naira | 2962.0 | 2987.1 | 3731.3 | 3731.3 | 0.450 | 6.6 | 8.7 | 83.0 |
| | 2024 | PC1 | 245.9 | 271.0 | 1015.2 | 1015.2 | 0.715 | 24.5 | 12.3 | 61.4 |
| | | PLER | 249.4 | 274.5 | 1018.7 | 1018.7 | 0.777 | 4.4 | 16.3 | 53.5 |
| | | Raw | 2774.6 | 2799.7 | 3543.9 | 3543.9 | 0.433 | 4.6 | 11.3 | 84.0 |
| | | Zscore | 431.0 | 456.1 | 1200.3 | 1200.3 | 0.715 | 24.5 | 12.3 | 61.4 |

AIC, Akaike information criterion; BIC, Bayesian information criterion; $H^2$, broad-sense heritability; GMA%, SMA%, Res%, percentage of total variance attributable to general mixing ability, specific mixing ability, and residual respectively. ΔAIC and ΔBIC are relative to the best-fitting metric within each trait-period combination.

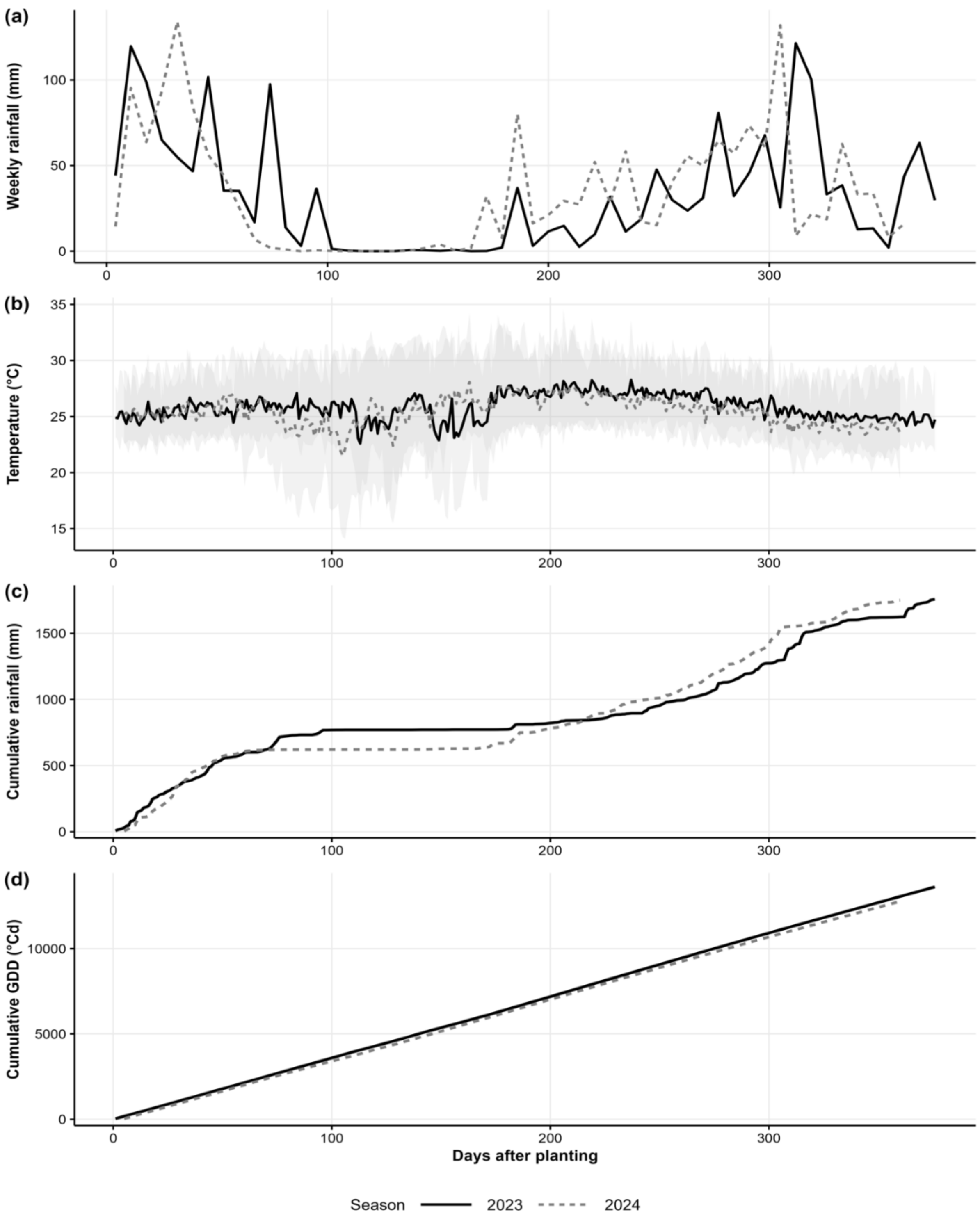


Supplementary Figure S1. Seasonal weather at the trial site

Figure S1. Weather across the two cassava–cowpea seasons (2023, solid; 2024, dashed) by days after planting: (a) weekly rainfall, (b) daily mean temperature (shaded band, daily minimum–maximum range), (c) cumulative rainfall, and (d) cumulative growing degree days (GDD). The seasons accumulated similar thermal time and total rainfall (~1,750 mm) but differed in rainfall distribution, with a pronounced mid-season dry spell in 2023 and a shorter low-rainfall window in 2024.

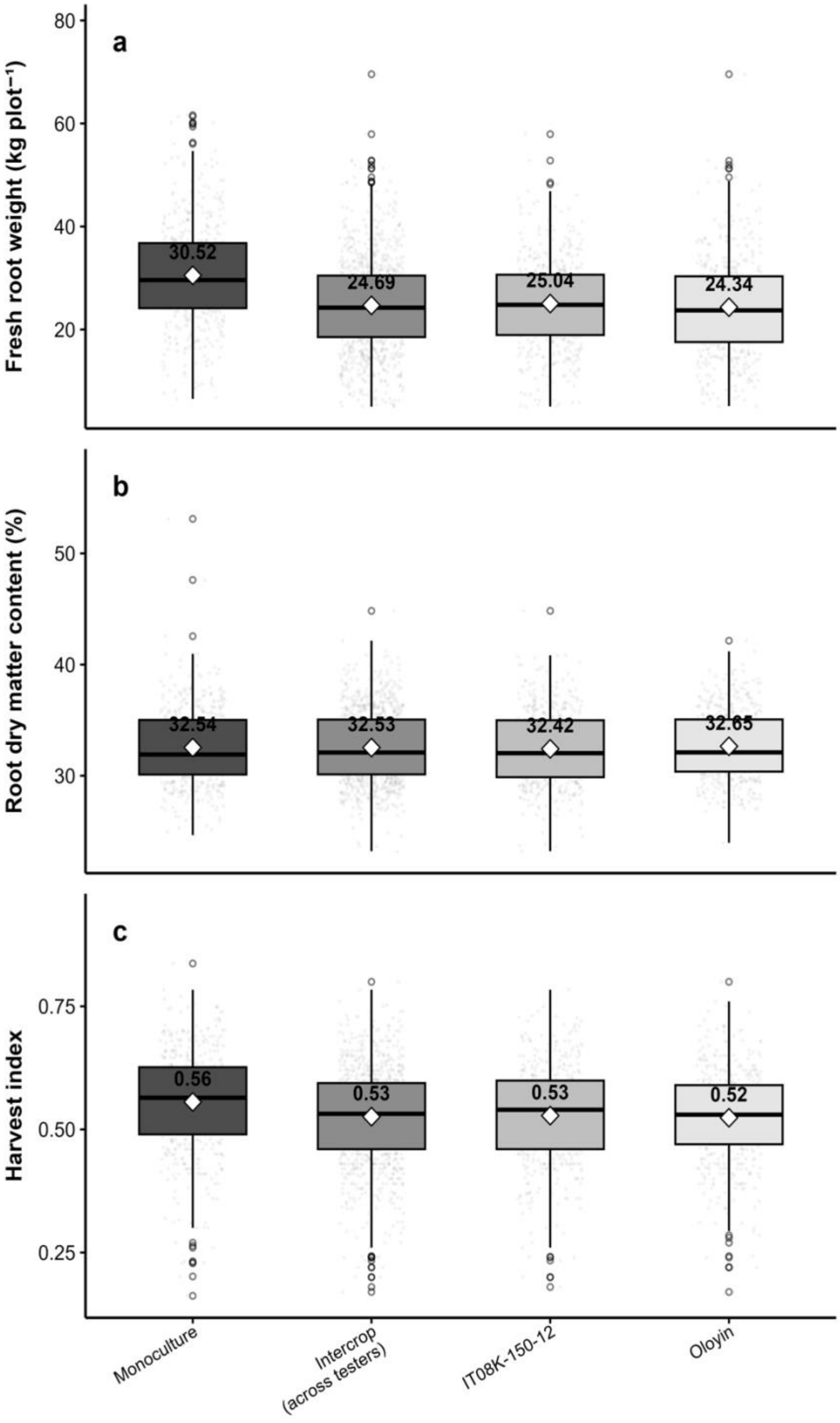


Supplementary Figure S2. Distribution of three cassava traits under monoculture and cassava cowpea intercropping, pooled across 2023 and 2024 at Ibadan, Nigeria

Figure S2. Distributions of three cassava traits, (a) fresh root weight, (b) root dry matter content, and (c) harvest index, under monoculture, intercrop (pooled across both cowpea testers), and each tester separately (IT08K-150-12, Oloyin), combined across 2023 and 2024. Boxes show the median and interquartile range, whiskers the 1.5× IQR, points individual plots, and white diamonds the means (labelled). Intercropping lowered fresh root weight (30.52 to 24.69 kg plot$^{-1}$) and slightly reduced harvest index (0.56 to 0.53), whereas dry matter content was essentially unchanged; differences between the two testers were small.

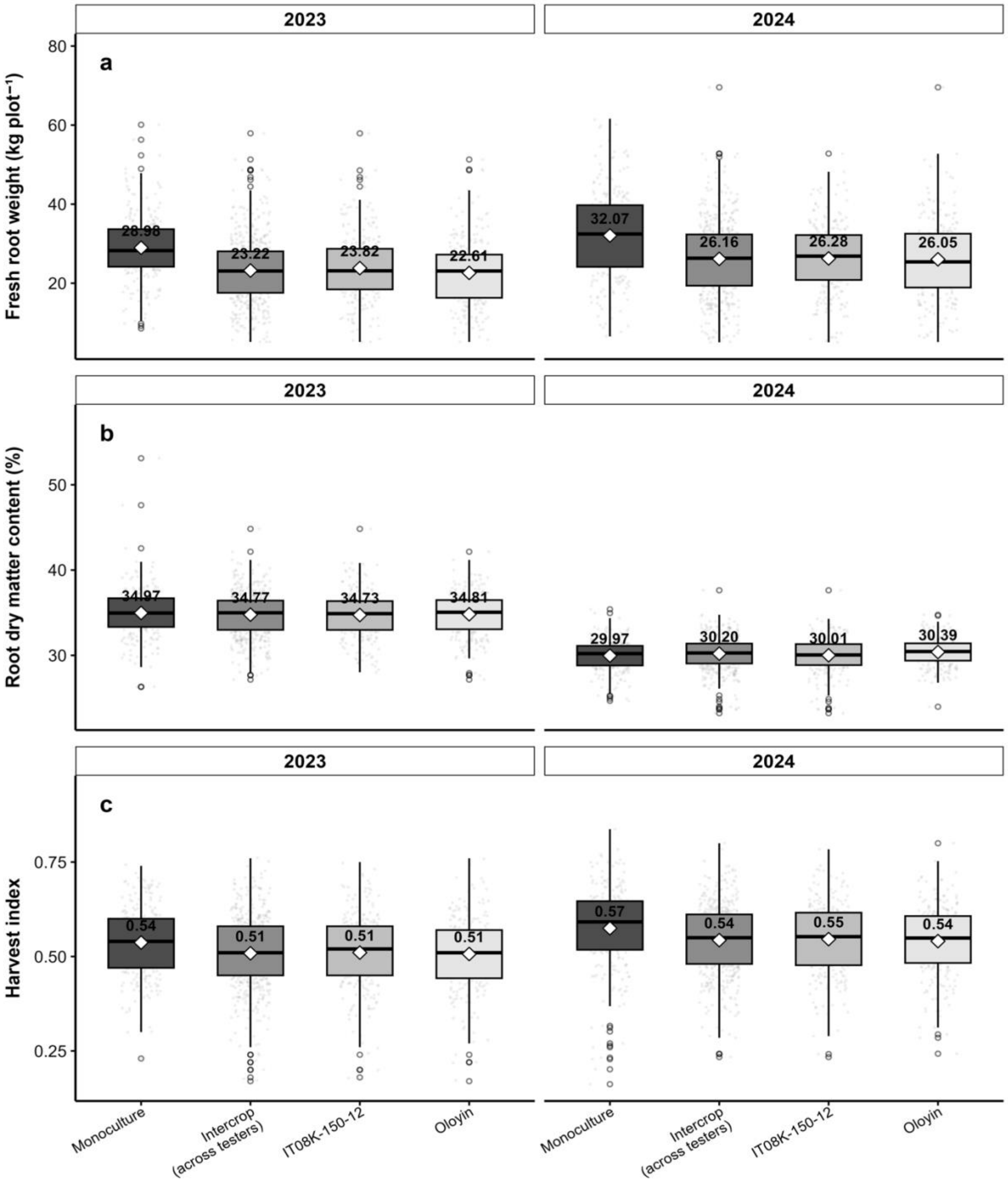


Supplementary Figure S3. Cassava trait distributions by cropping system and tester, within each year.

Figure S3. Distributions of three cassava traits, (a) fresh root weight, (b) root dry matter content, and (c) harvest index, under monoculture, intercrop (pooled across both cowpea testers), and each tester separately (IT08K-150-12, Oloyin), shown separately for 2023 (left) and 2024 (right). Boxes show the median and interquartile range, whiskers the 1.5× IQR, points individual plots, and white diamonds the means (labelled). The intercrop reduction in fresh root weight was consistent across years (28.98 to 23.22 kg plot$^{-1}$ in 2023; 32.07 to 26.16 in 2024), harvest index declined slightly, and dry matter content was largely unaffected by cropping system, though it differed markedly between years (~35% in 2023 vs ~30% in 2024). Tester differences were small in both years.

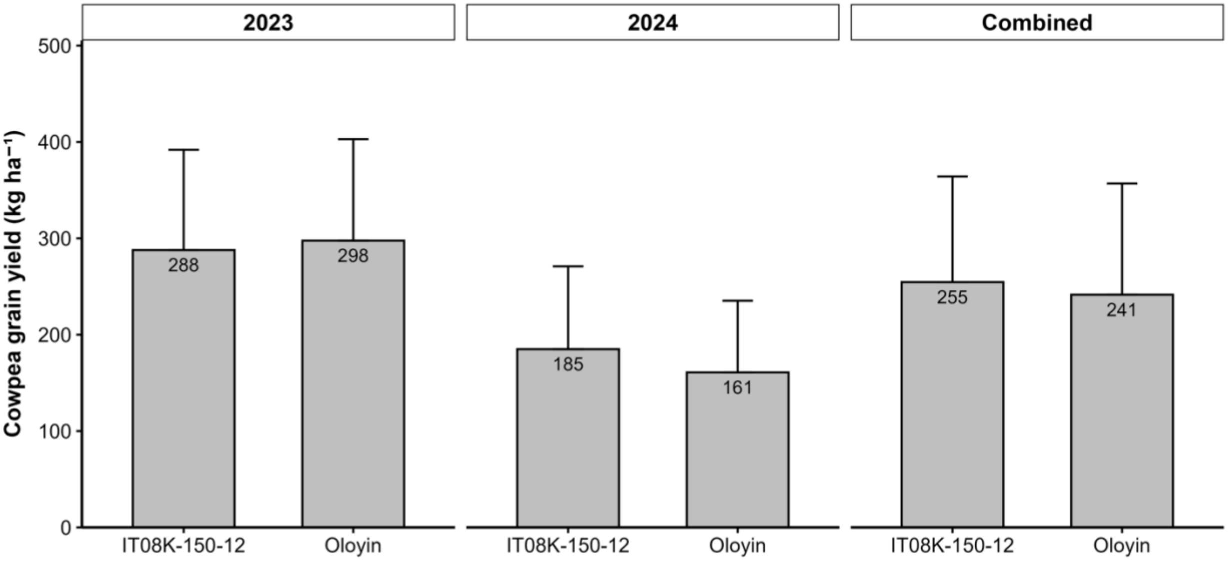


Supplementary Figure S4. Cowpea grain yield by tester
Figure S4. Intercrop cowpea grain yield (kg $ha^{-1}$) for the two testers, IT08K-150-12 and Oloyin, in 2023, 2024, and combined across years. Bars show means (labelled); error bars show the standard deviation. Grain yield was similar between the two testers in each panel but markedly lower in 2024 (185 and 161 kg $ha^{-1}$) than in 2023 (288 and 298 kg $ha^{-1}$), indicating that season influenced cowpea yield more strongly than tester identity.

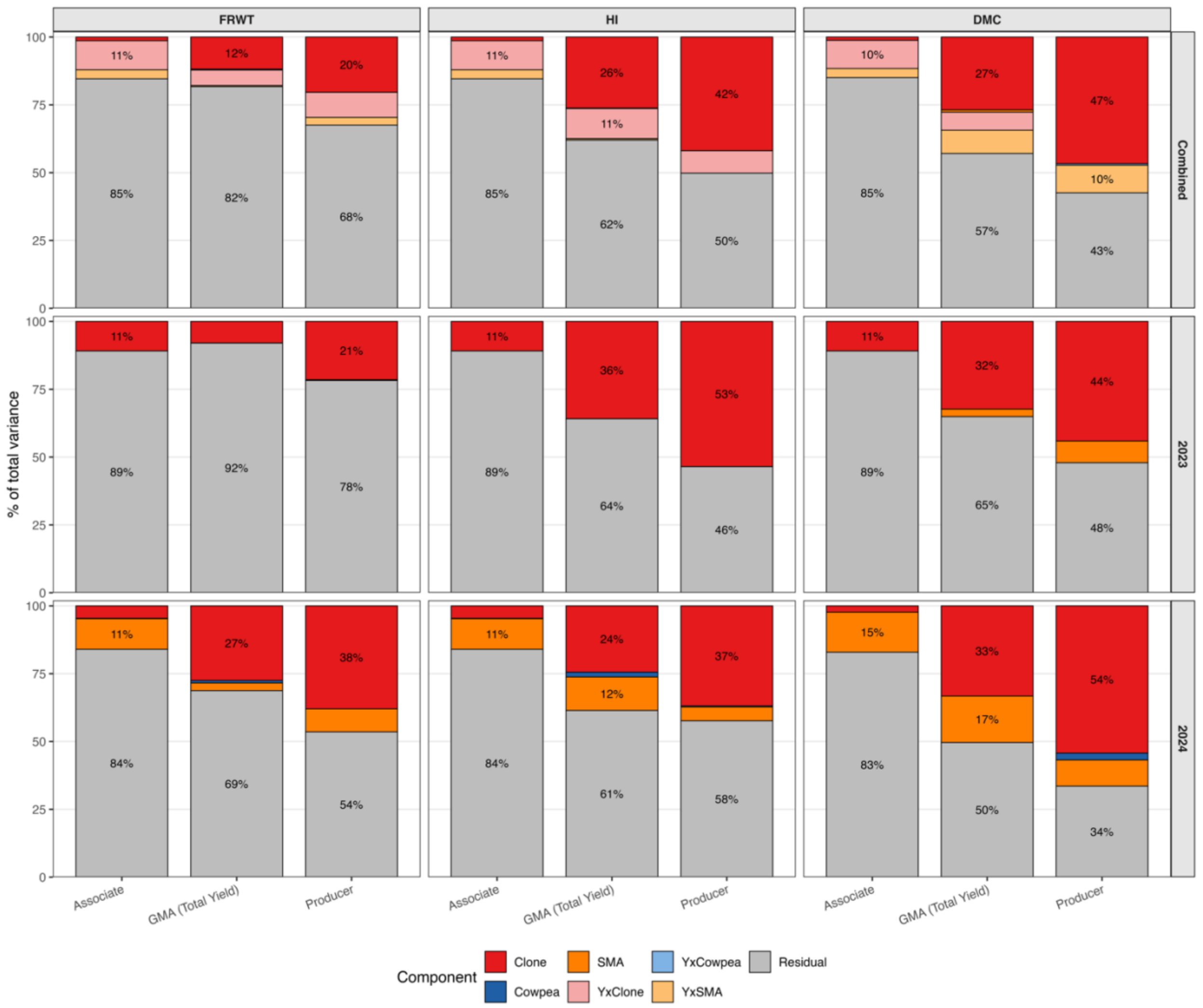


Supplementary Figure S5. Variance decomposition of the associate, general-mixing-ability, and producer models across traits and years.

Figure S5. Partitioning of total phenotypic variance under the associate, general-mixing-ability (GMA, total-yield merit index), and producer models, for fresh root weight (FRWT), harvest index (HI), and root dry matter content (DMC), in the combined-year (top), 2023 (middle), and 2024 (bottom) analyses. Each stacked bar shows the percentage of total variance attributable to clone, cowpea, specific mixing ability (SMA), the genotype-by-year interactions (Y×Clone, Y×Cowpea, Y×SMA), and residual; percentages are labelled for segments of 10% or more. Under the associate model the clone main effect was small (about 1% of variance) and the labelled segment near the top of each associate bar is the Y×Clone interaction; the model was otherwise residual-dominated. The clone component was largest under the producer model (up to 54% for DMC and 58% for HI in 2024) and modest under the GMA model, and clone contributions strengthened in 2024.

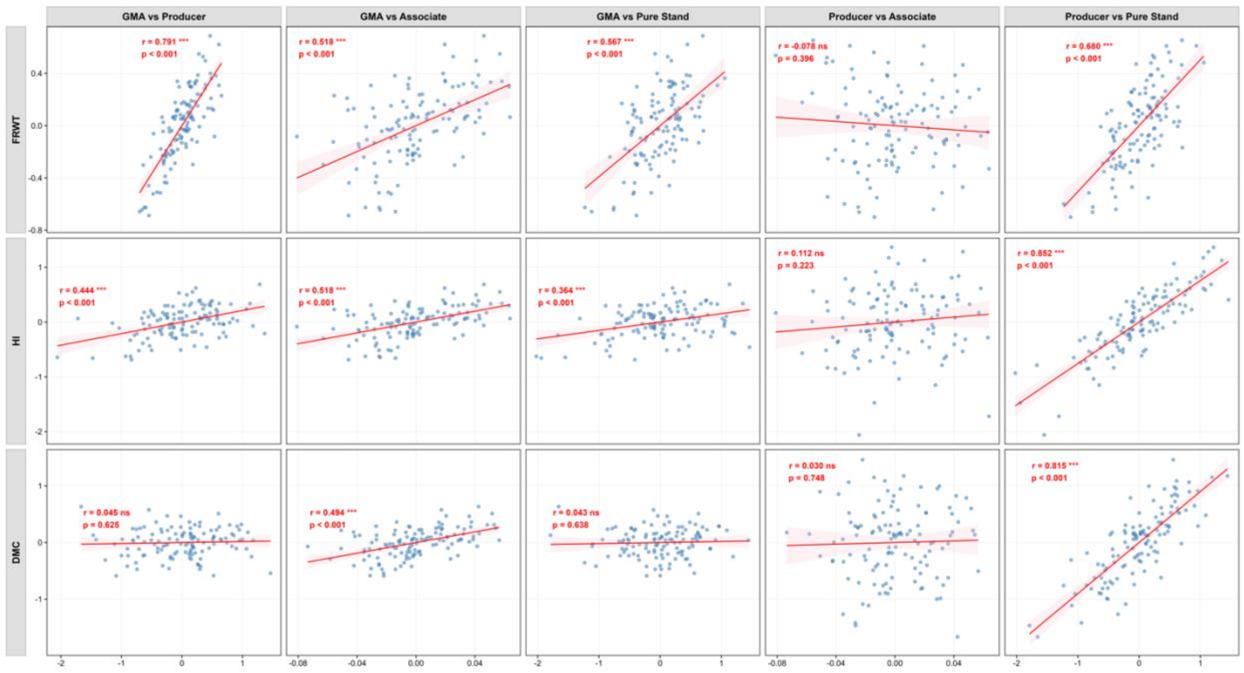


Supplementary Figure S6. Correlations among mixing-ability effects and monoculture performance

Figure S6. Pairwise relationships between clone BLUPs for general mixing ability (GMA), producer, associate, and monoculture (pure-stand) effects, for fresh root weight (FRWT), harvest index (HI), and root dry matter content (DMC), in the combined-year analysis. Each panel shows the Pearson correlation (*r*) with its significance and a fitted regression line (shaded 95% confidence band); columns are the five effect pairings and rows the three traits. GMA correlated positively with producer, associate, and monoculture effects, and producer effects correlated strongly with monoculture performance (FRWT *r* = 0.68, HI 0.85, DMC 0.82), whereas producer-to-associate correlations were weak and non-significant, indicating that monoculture selection captures the producer (direct) component but not associate effects. ns, not significant; *** $P < 0.001$.
